# Global error signal guides local optimization in mismatch calculation

**DOI:** 10.1101/2025.07.07.663505

**Authors:** John Hongyu Meng, Xiao-Jing Wang

## Abstract

Corollary discharge denotes internal signals about the expected sensory consequences of one’s own actions, leading to attenuation of sensory responses caused by self-produced stimulation. To investigate the underlying neural circuit mechanism, here we introduce a biologically plausible three-factor learning rule, where a global signal guides the updating of local inhibitory synapses to enable the computation of mismatch between a stimulus and its expectation or prediction. We show that our network model, endowed with positive and negative prediction error neurons, accounts for the salient physiological observations of motor-visual and motor-auditory mismatch responses in mice. Moreover, the model predicts that learning induces a bimodal distribution in activity correlation with stimulus and movement-induced prediction, supported by our analysis of neural data from a recent experiment. These results link global modulation to local learning for predictive error computation in the sensory areas, and shed insights into how disrupting inhibition impairs mismatch computation in specific ways.

## 2 Introduction

Since Hermann von Helmholtz, an outstanding question in Psychology and Neuroscience has been to elucidate how self-generated movement is monitored in the brain through an internal signal called corollary discharge or efference copy, that conveys (top-down) expectation of sensory consequences of one’s own actions, to be compared with the actual stimulus induced (bottom-up) input in a sensory system [15] (Figure 1A). For instance, we make rapid eye movements several times per second. Yet, our vision is stable thanks to corollary discharge associated with saccades [55]. When we walk, we do not perceive that the world is moving, even though the visual input to our eyes suggests motion [38]. Similarly, when playing an instrument in a band, we may tune out the sound of our own actions relative to those of others [9]. What neural mechanisms underlie this selective suppression? Answering this question is of interest for basic neuroscience as well as psychiatry, since deficits in corollary discharges are believed to be a root cause of hallucination and delusion associated with mental disorders [22, 13]. In recent years, mismatch computation between a sensory response and corollary discharge of self-motion has been cast in the general framework of predictive coding [34]. Originally proposed for perception, predictive coding highlights the idea that a stimulus-induced response in a sensory circuit is compared with an internally generated expectation/prediction signal about that stimulus, and is suppressed when expectation is increased [47]. Applied to motor-sensory feedback, the motor cortexsends a copy of the movement signal to sensory areas, which serves as a local prediction of the incoming stimulus. If the prediction and actual input match, their signals cancel out, leading to reduced or no sensory perception (Figure 1B). For example, we may press a play button and hear the expected tune. But when the prediction and stimulus mismatch, the sensory area responds, signaling a prediction error (Figure 1C). This mismatch can occur in two ways: either a sound is heard without pressing the button, or we press the button while no sound follows. Both types of mismatch elicit stronger neural responses in mice than expected stimuli [2, 4]. Moreover, suppression in the expected case depends on experience, suggesting a critical role for neural plasticity.

**Fig. 1.**
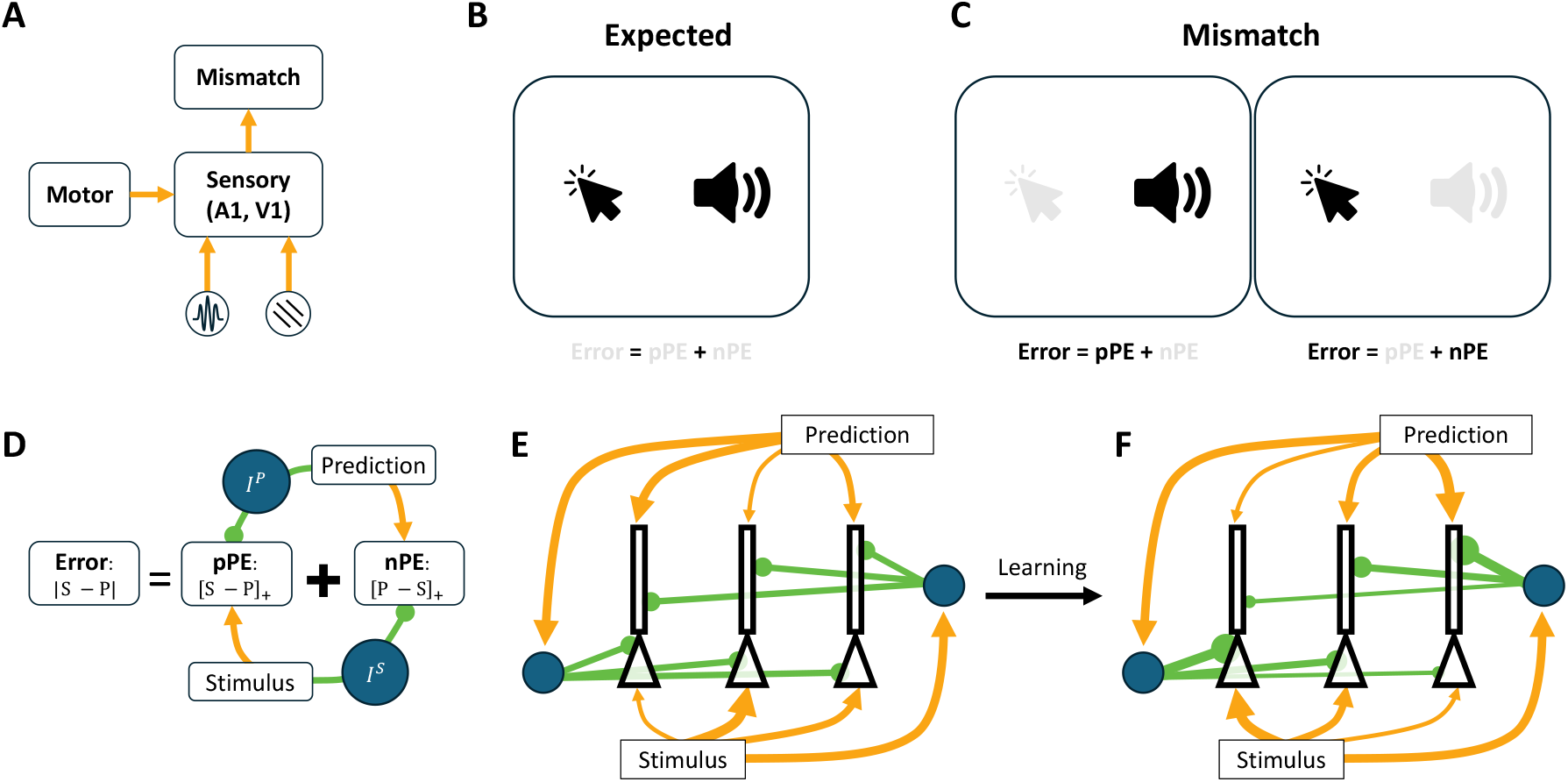
Motion-sensory mismatch calculation requires learning. (A) An efference copy of the motion signal is compared with the incoming auditory or visual stimulus to compute the mismatch. (B) A matched case where the incoming sound aligns with the clicking motion. (C) Mismatch examples: (left) sound is heard without a corresponding motion, or (right) no sound is heard when clicking the button. (D) The mathematical description of error calculation requires two distinct prediction error populations and local interneurons to invert long-range excitation into local inhibition (see main text). (E) Innate connectivity fails to compute errors effectively. (F) Optimal connectivity for error calculation requires inhibitory input from prediction (stimulus) to balance excitation from stimulus (prediction), a pattern that can be achieved through learning.

Though the mathematical description of mismatch is as simple as computing the absolute difference between stimulus and prediction signals, it is not clear how the brain implements this operation (Figure 1D). One challenge is that neurons in sensory areas exhibit low baseline firing rates [16, 42, 49], making them highly responsive to excitatory input but relatively insensitive to inhibition. A straight-forward solution is to introduce two separate neuronal populations: positive prediction error (pPE) neurons, which respond to stimulus minus prediction, and negative prediction error (nPE) neurons, which respond to prediction minus stimulus [34]. Another challenge is that long-range projections in the brain are predominantly excitatory. Therefore, to compute a difference between stimulus and prediction signals, local interneurons must be involved to convert long-range excitatory inputs into local inhibition.

However, accurate prediction error computation also requires well-tuned local connectivity. With naive connectivity (Figure 1E), stimulus and motor-related inputs can vary across individual neurons (Figure 1D). Yet, these neurons receive dense inhibition from local interneuron populations [33], suggesting that each neuron receives a comparable level of inhibition. As a result, expected stimulus input cannot be effectively cancelled by prediction-driven inhibition, and vice versa. This mismatch implies the need for a learning mechanism that tunes local inhibitory connections to accurately align inhibition with its corresponding excitatory drive, such that relayed stimulus inhibition targets prediction-driven excitation and relayed prediction inhibition targets stimulus-driven excitation. (Figure 1E). Notably, excitatory plasticity may not play a major role, as overall excitatory responses to motion or visual input are not significantly altered after learning [2]. Taken together, successful prediction error computation likely depends on inhibitory plasticity. This form of plasticity has been widely observed in cortex [60, 36, 11] and is modulated by neuromodulators [10]. In particular, noradrenaline signals broadcast by the locus coeruleus may facilitate local circuit optimization [32, 30], though the underlying mechanism remains unknown.

In this study, we introduce a biologically plausible three-factor learning rule, inspired by [24], that acts on inhibitory synapses from interneurons to pyramidal cells and enables local circuits to compute prediction errors. We show that this local rule converges to the same optimal connectivity as that learned via gradient descent. We test the rule in models composed of either simplified ReLU or biophysically realistic neuron models. The resulting inhibitory connectivity reproduces key experimental observations, including the selective motion-modulated auditory response [2] and the motion-visual mismatch response [4]. Moreover, we demonstrate that this learned connectivity is functionally optimal: perturbing interneuron activity, either by excitation or inhibition, consistently disrupts the population-level mismatch response. Finally, the model predicts a bimodal distribution of stimulus- and prediction-related correlation across neurons, a feature supported by reanalysis of experimental data.

## 3 Results

### 3.1 Three-factor learning rule achieves the optimal solution in computing prediction error

We implemented a biologically feasible three-factor learning rule to align inhibition relayed from the prediction (or stimulus) with excitation driven by the stimulus (or prediction), such that population responses are suppressed when inputs are predicted, but enhanced during mismatches. This learning rule depends on the firing rates of the pre- and postsynaptic neurons, along with a global third factor representing the mismatch between prediction and stimulus. Inspired by recent studies [32, 30], this third factor may be mediated by noradrenaline, released from the locus coeruleus. Remarkably, the proposed rule converges to the same optimal solution as the traditional gradient descent algorithm, despite relying only on biologically plausible global error signals. The detailed mathematical equivalence is provided in the Supplementary Material. Here, we offer an intuitive explanation.

We began with a simplified case of *n* = 2 rectified linear units (ReLU), consisting of one pPE and one nPE neuron (Figure 2A), each projecting equally to an output neuron whose activity encodes the error between stimulus and prediction. In this simplified case, we also assume that the stimulus input is exclusively to the pPE neuron and the prediction input is exclusively to the nPE neuron. We then trained the inhibitory connections from two interneurons *I*^*S*^, *I*^*P*^ to both the pPE and nPE neurons to minimize the difference between the output neuron’s activity and the true error, defined as the absolute difference between the stimulus and prediction. The optimal solution in this setting requires the connectivity configured as in Figure 1D. In this case, the pPE neuron only receives relayed prediction inhibition but none of the relayed stimulus inhibition, and its firing rate reflects the positive prediction error *R*^pPE^ = [*S* − *P*]_+_. Conversely, the nPE neuron only receives relayed stimulus inhibition but not prediction inhibition, and its firing rate reflects the negative prediction error *R*^nPE^ = [*P* − *S*]_+_.

**Fig. 2.**
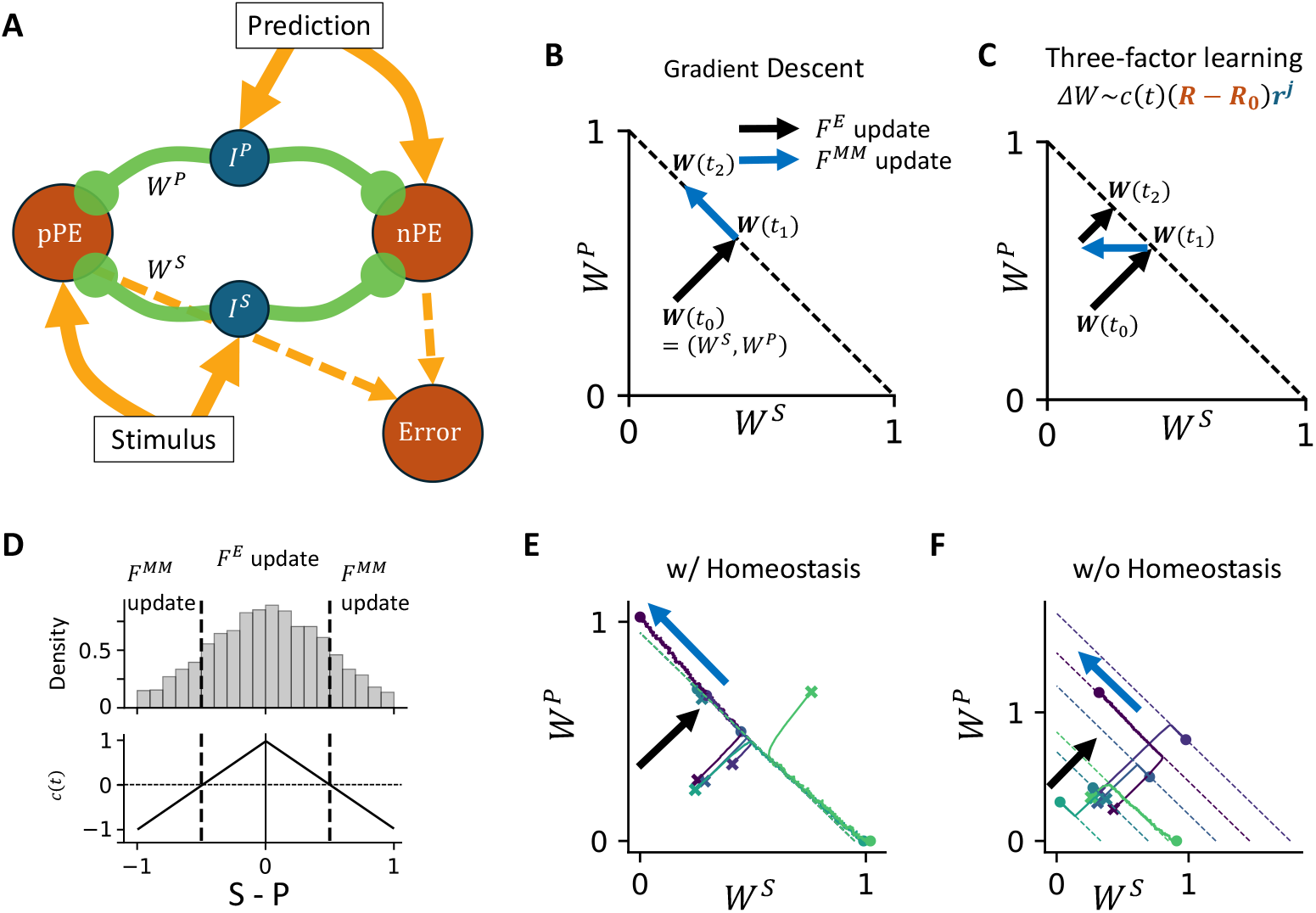
Mechanisms of the three-factor learning rule. (A) Diagram of network connectivity in a simplified case with *n* = 2, consisting of one nPE and one pPE neuron. (B) Weight updates under gradient descent. When trained on the expected set *F* ^*E*^, where prediction matches stimulus, weights converge to the slow learning manifold *W* ^*S*^ + *W* ^*P*^ = 1. Constrained by the slow learning manifold, training on the mismatch set *F* ^*MM*^ shifts the weights along it. Eventually, inhibitory selectivity emerges as (*W* ^*S*^, *W* ^*P*^) *→* (0, 1). (C) Weight updates under the local three-factor learning rule. Inhibitory selectivity emerges through a zig-zag trajectory guided by the local rule. *c*(*t*) represents the third factor. See Methods for details. (D) The sign of the third factor *c*(*t*) depends on the true error. When the mismatch is small (|S − P |*<* 0.5), weights update as in the expected set *F* ^*E*^; when large (|S − P |*>* 0.5), weights update as in the mismatch set *F* ^*MM*^. (Top) The error distribution is drawn from a truncated Gaussian distribution. (Bottom) Corresponding third factor *c*(*t*). (E) Weight evolution when total excitatory input from stimulus and prediction is the same across neurons (homeostasis hypothesis). The dashed line indicates the theoretical slow-learning manifold. Crosses mark initial weights; circles mark final weights; colors mark different post-synaptic neurons. The black arrow highlights fast initial learning; the blue arrow indicates slower refinement along a slow learning manifold. (F) Same as (E), but without the homeostasis hypothesis. In this case, the total inhibitory input diverges across neurons.

Learning how to compute prediction error accurately requires a model or an animal to use the prediction signal to cancel the incoming stimulus in the expected cases when both the prediction and stimulus signals are present (*F*^*E*^ : |*S* − *P* | *≈* 0). However, the optimal configuration cannot be reached only by training in the expected case. This can be illustrated by examining the behavior of updating the synaptic weights through the gradient descent algorithm. Within the expected training set where stimulus and prediction inputs are matched (*F*^*E*^ : |*S* − *P* | *≈* 0), the total inhibitory input converges to match the total excitation (Figure 2B, black arrow). Yet, in this condition, it does not matter whether the inhibition originates from the prediction or the stimulus, resulting in a family of equally valid solutions rather than a unique one. This family of solutions is referred to as the slow learning manifold in the following. Indeed, the optimal response in the expected case is zero, indicating that excessive inhibition can trivially suppress neural activity, thereby satisfying the objective without yielding a meaningful solution. To resolve this ambiguity, training on the mismatch set *F*^*MM*^ : |*S* − *P*| *>* 0, where stimulus and prediction inputs are not matched, is necessary. In this case, the source of inhibition becomes critical for accurate mismatch computation. As a result, the synaptic weights move along the slow learning manifold to the unique optimal solution (Figure 2B, blue arrow), where the circuit correctly associates inhibition with its corresponding excitatory drive.

The same optimal solution emerges from our three-factor learning rule, assuming expected training samples are more frequent than mismatched ones, as is typical in experimental paradigms. The third factor, *c*(*t*), presumably mediated by a neuromodulator such as noradrenaline [32], controls the sign of learning. Indeed, [53] demonstrated that different concentrations of neuromodulators could control whether paired pre- and post-synaptic activity led to long-term potentiation or depression. During training on expected samples, the rule behaves like Hebbian learning, with weight updates proportional to the product of pre- and post-synaptic activity, causing total inhibition to match excitation and thereby defining a slow-learning manifold. For mismatched samples, the rule becomes anti-Hebbian, with weight updates negatively correlated with the product of activity, driving weights toward the optimal configuration. Although inhibition temporarily deviates from the slow-learning manifold, the dominance of expected samples pulls total inhibition toward the manifold, keeping it close throughout learning. This alternation guides convergence along a zigzag trajectory (Figure 2C).

Next, we test our algorithm with a network of *n* = 40 ReLU neurons. At each learning step, the true prediction error is sampled from a truncated Gaussian distribution (Figure 2D), and the third factor *c*(*t*) reflects the true error. Here, the third factor is only required to differentiate expected cases from mismatched ones by its sign, while the exact form may vary. Here, we choose a linear function for simplicity. For analytical convenience, we assume that the sum of stimulus and prediction inputs is constant across neurons, a condition we refer to as the homeostasis hypothesis [48, 19]. Under this assumption, different neurons share the same slow learning manifold, which is defined by the total excitatory input, and they can be ordered along a one-dimensional axis according to their input affinity to the stimulus (Figure 2E). We further assume that the utility of mismatch signals saturates beyond a certain threshold (see Methods), which causes synaptic weights to different postsynaptic neurons to converge at distinct positions along the slow learning manifold. However, the homeostasis assumption may not hold in cases where a neuron responds to multiple stimuli with varying strengths. Indeed, homeostasis is not required for learning in our model, where the slow learning manifold differs for each neuron (Figure 2F). The corresponding activity and synaptic weights across neurons are shown in Supplementary Figure 9.

### 3.2 Prediction response depends on context after learning

After establishing our learning rule, we examine neural responses to stimulus and prediction signals before and after learning. From this point on, we use more realistic models for both pyramidal neurons and interneurons (see Methods), though ReLU neurons yield similar results (Supplementary Figure 9). Our pyramidal neuron model consists of two compartments and includes a dendritic voltage variable. With weak coupling between the somatic and dendritic compartments, the model captures both subtractive and divisive dendritic inhibition, depending on the level of dendritic depolarization (Supplementary Figure 10). Here, we use the model with strong dendro-somatic coupling, reflecting a model of Layer 2/3 pyramidal neurons. We further select the single-neuron parameters based on data recorded from Layer 2/3 in mouse primary visual cortex [25]. Neurons receive a baseline input that maintains low but nonzero firing rates. Each neuron receives input from both the stimulus and prediction signals, with varying strengths. They also receive inhibition that is relayed through local interneuron populations. Given the high density of local inhibitory connectivity [33], we assume that the strength of inhibition is similar across neurons. For simplicity, there is no recurrent connectivity among pyramidal neurons or from pyramidal neurons to interneurons in this model.

Before learning, the majority of neurons exhibit stronger responses when both the stimulus and prediction are present, compared to when only one input is provided (Figure 3A). In this condition, each neuron receives the same total input in the expected case (the homeostasis hypothesis), and neurons are sorted based on their affinity to the stimulus. At the population level, the response is highest when both inputs are present. Under our learning rule, inhibitory connectivity evolves from uniform inhibition into selective inhibition, such that excitatory input from the stimulus is cancelled by inhibition relayed from the prediction (Figure 3B, top), and excitatory input from the prediction is cancelled by inhibition relayed from the stimulus (Figure 3B, bottom).

**Fig. 3.**
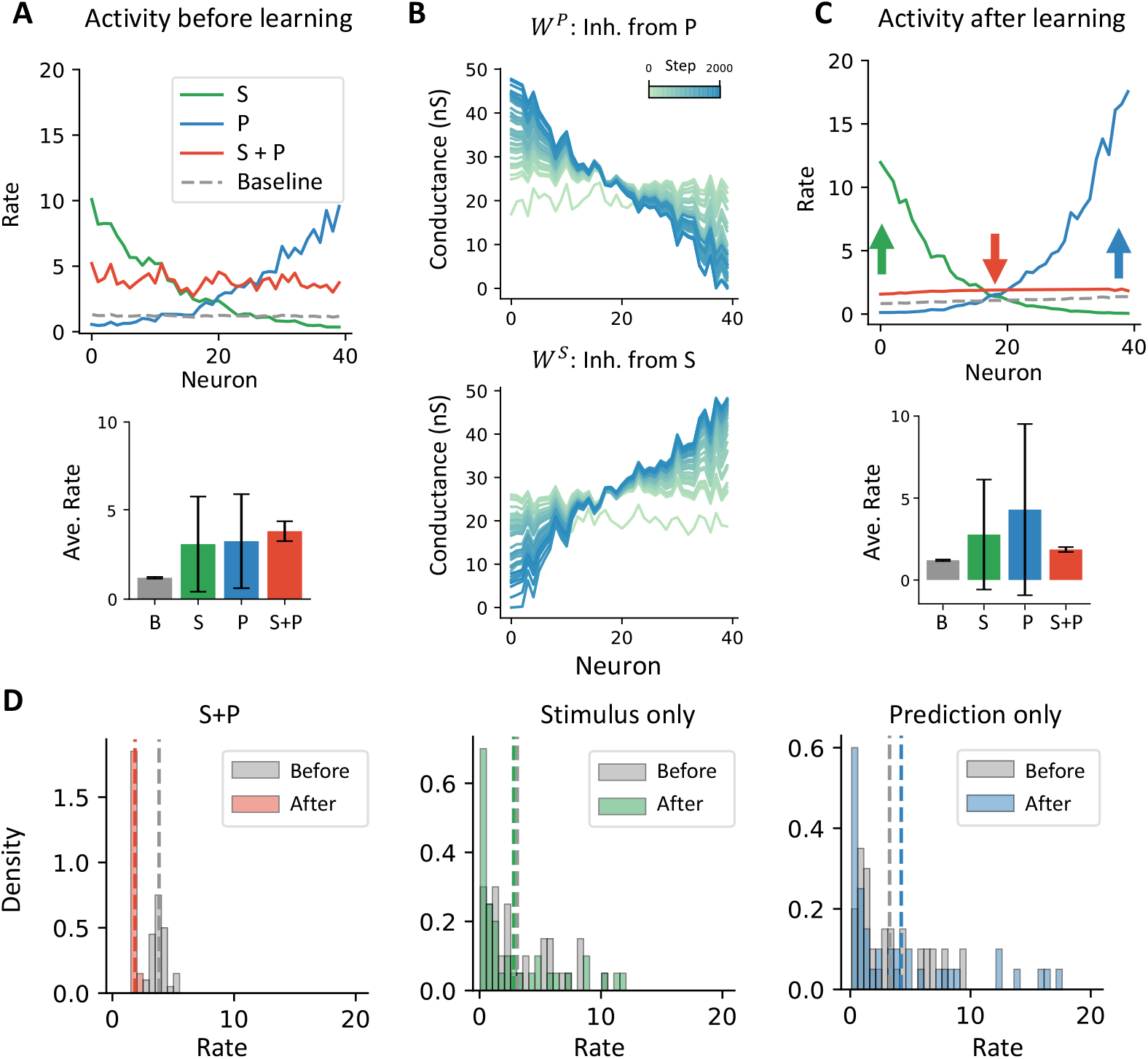
Neuron activity before and after learning. (A) Neuron activity before learning. (Top) The activity of individual neurons is sorted by the stimulus input strength. The line color indicates the type of input: Only stimulus, only prediction, stimulus and prediction, or neither (baseline). (Bottom) The bar chart of the average rate. The error bar indicates the standard deviation. (B) Connectivity weight that relays prediction inhibition *W* ^*P*^ (Top), and that relays inhibition from stimulus inhibition *W* ^*S*^ (Bottom). Different colors represent synaptic weight during different training steps. A darker color indicates a later connectivity in the learning. (C) Neuron activity after learning. The activity is minimal in the expected case when both stimulus and prediction input are received, compared to the mismatched cases, when only stimulus or only prediction input is presented. (D) Activity distribution before and after learning across different inputs. Population activity only decreases when both prediction and stimulus are simultaneously presented.

After learning, population neuronal activity reflects prediction error computation (Figure 3C). The left-most neuron receives the strongest stimulus excitation and the least inhibition from the stimulus, resulting in stronger responses when only the stimulus is present. The right-most neuron shows a similar pattern when only the prediction input is provided. However, when both stimulus and prediction inputs are present, all neurons exhibit low firing rates, slightly above baseline. At the population level, the average response in the expected condition, when both stimulus and prediction inputs are present, is lower than the response to either input alone. Furthermore, comparing activity distributions before and after learning, the average activity decreases only in the expected condition (Figure 3D). The model has similar qualitative results without the homeostasis assumption (Supp. Figure 11).

From another perspective, the net effect of the prediction signal on population responses becomes context-dependent after learning. Before learning, adding prediction increases activity, both with the stimulus (Figure 3A, blue vs. grey) and without it (Figure 3A, red vs. green). After learning, however, prediction reduces activity when comparing the expected case to the stimulus-only case (Figure 3C, red vs. green), indicating that prediction-modulated inhibition is learned and context-specific. We will examine this further in the next section.

Here, the exact form of the third factor *c*(*t*) is not important. To directly test this, we use a piecewise-estimated 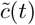 instead to guide the learning and reach the same qualitative results (Supp. Figure 12). In this case, the 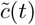 requires only an approximation of the global error by the animal.

Since the neurons on the left respond selectively to positive prediction error, [S − P]_+_, after learning, we refer to them as pPE neurons. The same applies to the neurons on the right, which respond to negative prediction error, and are referred to as nPE neurons. Along the x-axis of input strength, the neuron identity gradually transitions from pPE (neuron 1 to neuron 20) to nPE (neuron 21 to neuron 40). In the following sections, the representative pPE and nPE neurons correspond to neuron 1 and neuron 40, respectively.

### 3.3 Prediction input selectively modulates the expected stimulus

In many situations, we selectively suppress expected stimuli in complex environments, while leaving the representation of other stimuli unchanged. Recent experiments have shown that this suppression is highly specific: only the response to the expected sound associated with self-generated motion is reduced, whereas responses to other, non-predicted sounds remain unaffected [4, 3]. Does the prediction error computation learned by our model show the same specificity to expected stimuli?

We examine this selectivity by introducing a probed stimulus in addition to the expected stimulus. The probed stimulus is generated by shuffling the expected stimulus. In addition, we do not assume homeostasis here since the response to two stimuli is of interest. We do so by shuffling the prediction input to eliminate any correlation with either the expected stimulus or the probed stimulus (Figure 4A).

**Fig. 4.**
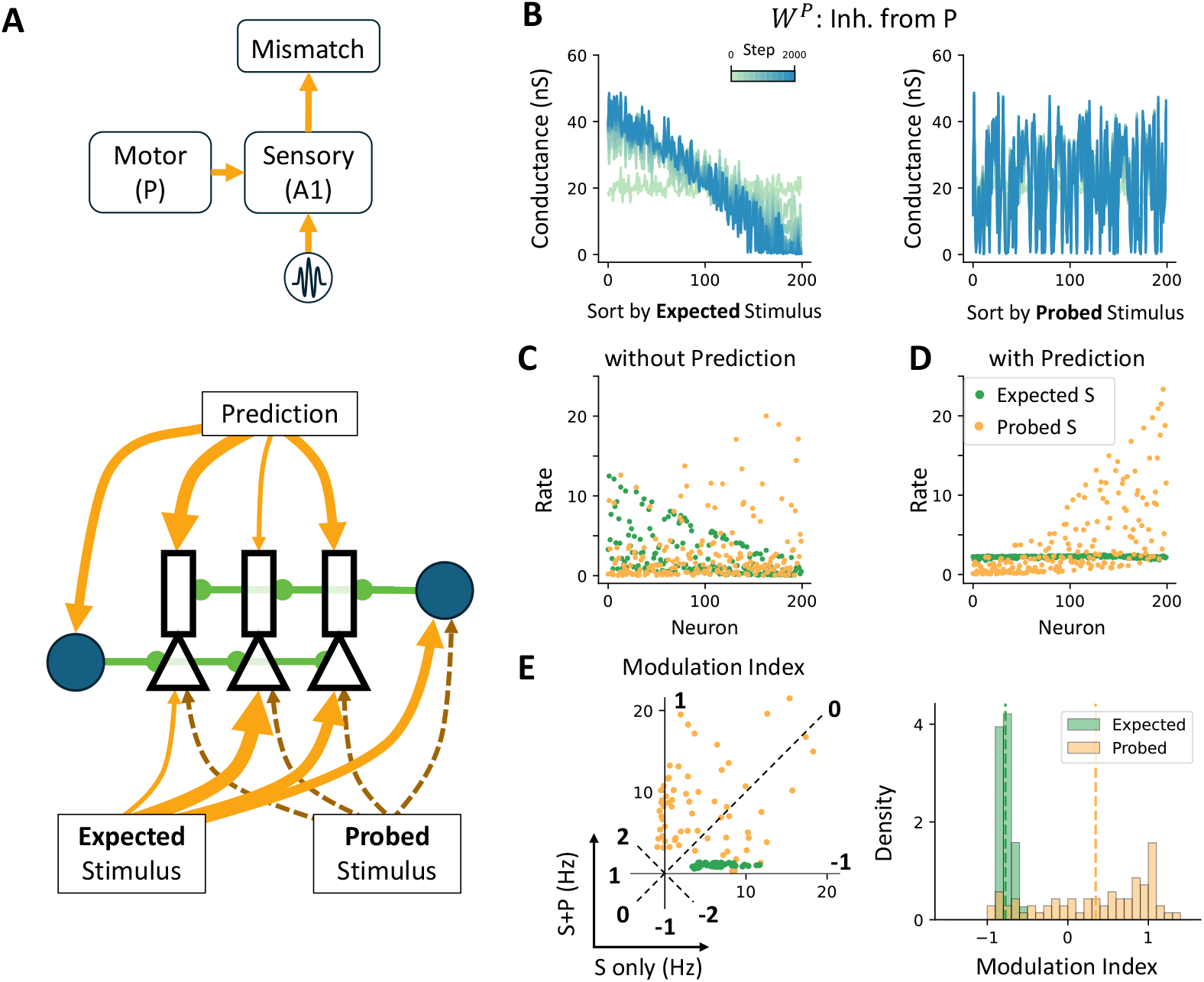
Selective suppression in the learned connectivity. (A) Schematic of stimulus-specific suppression. (Top) A1 compares motor and auditory inputs to generate a stimulus-specific mismatch signal. (Bottom) Schematic of the model setup. During learning, an expected stimulus is presented. A probed stimulus is generated by shuffling the original stimulus inputs across neurons. (B) Learned inhibitory connectivity (*W* ^*P*^) aligns with the expected stimulus. (Left) *W* ^*P*^ is strongly aligned with neurons receiving stronger input from the expected stimulus. (Right) No such alignment is observed when neurons are sorted by probed stimulus input. (C) Firing rates in response to expected or probed stimuli alone. Neuronal activity does not differ significantly between the two conditions when no prediction input is present. (D) Selective suppression emerges when prediction is added. The expected stimulus is selectively inhibited in the presence of its associated prediction, whereas the probed stimulus is not. (E) Quantifying suppression with a modulation index. (Left) A scatter plot of responses to stimulus alone (x-axis) vs. stimulus with prediction (y-axis) shows stronger suppression for the expected stimulus. (Right) Distribution of the modulation index shows a peak near −1 for the expected stimulus, consistent with strong learned suppression. (E) and (F) agrees with the selective suppression observed in [4, 3].

During training, as shown previously, the synaptic weights gradually align the inhibition relayed from the stimulus with the prediction input (Supplementary Figure 13A). In contrast, inhibition relayed from the prediction aligns specifically with the expected stimulus, but not with the probed stimulus (Figure 4B). This is because the error signal driving the third-factor learning rule reflects only the mismatch between the expected stimulus and the prediction, and does not incorporate the probed stimulus.

After learning, in the absence of prediction input, the response distributions to the expected and probed stimuli are similar (Figure 4C). In contrast, when both stimulus and prediction inputs are present, strong inhibition emerges selectively for the expected stimulus, but not the probed stimulus (Figure 4D, Supplementary Figure 13B).

To quantify the prediction-modulated response, we used the same modulation index (MI; see Methods) as in [4, 3] to measure activity changes within the same neuron population (Figure 4E). The MI for most neurons for the expected stimulus is close to minus one, indicating strong selective inhibition. In contrast, the MI for the probed stimulus shows a broader distribution centered above zero, consistent with weaker or absent modulation. This difference in MI between expected and probed stimuli was also observed experimentally [4, 3]. One notable difference is that, in the experiments, the MI for the probed stimulus is closer to zero or slightly negative. This discrepancy may be explained by a global, non-specific inhibitory effect from motor to auditory cortex, potentially mediated by Parvalbumin-expressing (PV) interneurons [4].

It is also important for the sensory cortex to maintain accurate stimulus processing in the absence of prediction input. We find that discriminability, measured using a form of the Fisher information metric (see Methods), is comparable between the expected and probed stimuli (Supplementary Figure 13C).

### 3.4 Model reproduces motor-visual mismatch response after learning

In the previous section, we examined how neural responses depended on whether a stimulus was predicted. Here, we investigate a complementary scenario in which the stimulus is absent when a motor signal is present (Figure 5A). Recent studies [2, 32] used a motor-visual mismatch protocol to explore neural responses when expected sensory input was omitted. In this paradigm, dark-reared mice were passively trained in a virtual environment where their movement on a treadmill controlled the visual flow of a grating pattern (coupled training, CT). In parallel, a control group of mice received the same visual input generated from the CT animals, but the visual flow was decoupled from their own movement (non-coupled training, NT). As a result, the CT group may learn that the visual flow was a consequence of their movement, but not the NT group. After training, Mismatch events were introduced by abruptly pausing the visual flow during locomotion for both groups. In these experiments, a larger fraction of neurons in the CT group responded to the mismatch, and the average mismatch response was significantly stronger in the CT group compared to the NT group.

**Fig. 5.**
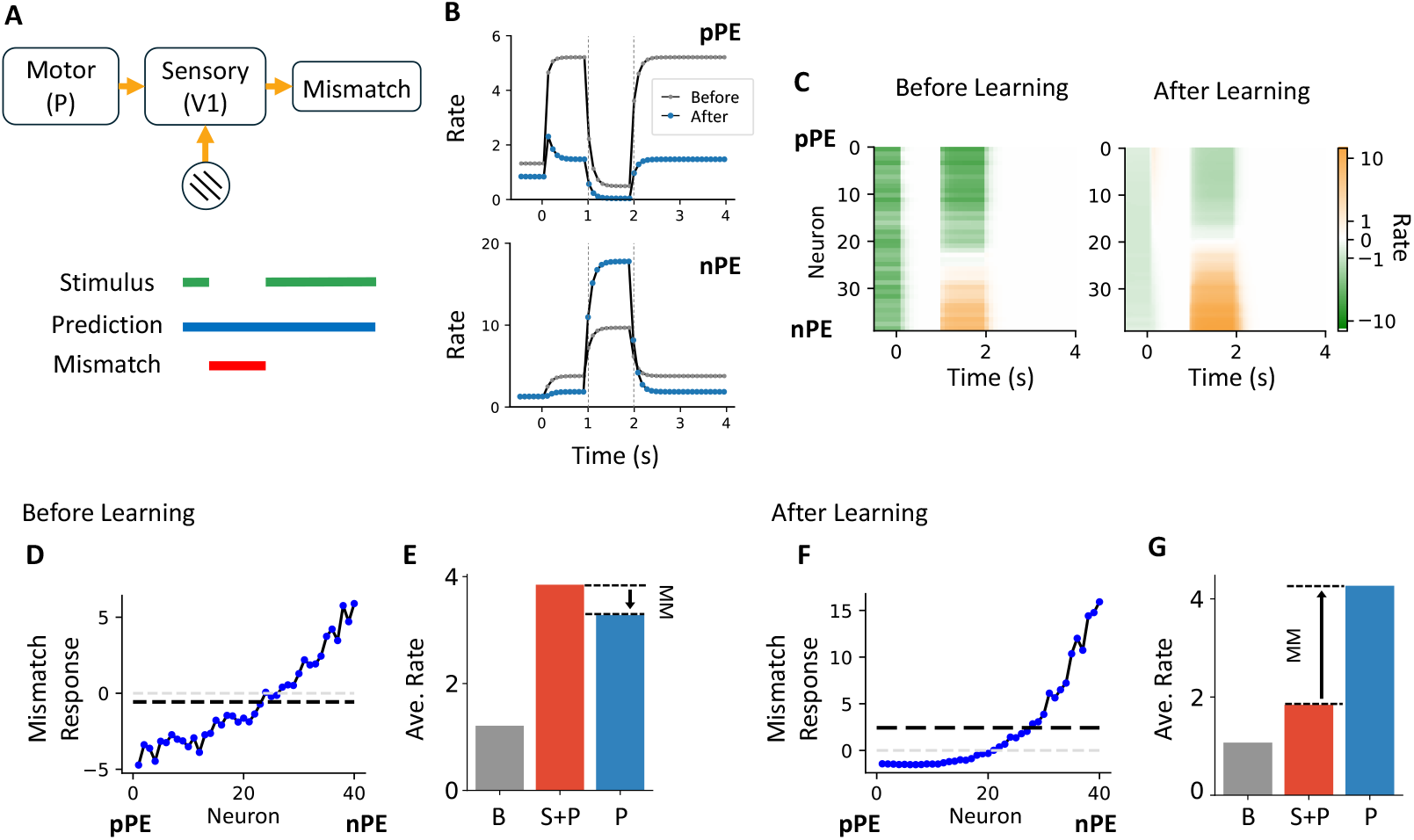
Three-factor learning captures responses in a motor-visual mismatch protocol. (A) Schematic of the mismatch paradigm. (Top) V1 compares motor-related and visual inputs to generate a surprise signal. (Bottom) Task schematic: A mismatch is introduced by pausing the flow of visual stimuli during locomotion. (B) Example neuron responses before and after learning. Firing rates over time for one pPE and one nPE neuron are shown before learning (black) and after learning (blue). (C) Learning-induced changes in firing rates. Firing rate changes across all neurons, comparing pre- and post-learning conditions. (D) Mismatch responses before learning across neurons. The black dashed line indicates the population average. (E) Population activity before learning. The averaged response in the expected case (S+P) is higher than the mismatch case (P only), suggesting no mismatch response at the population level. (F, G) Mismatch responses after learning. Same as (D, E). Here, a mismatch response is detected at the population level. The model behavior before learning and after learning reflects the behavior of the NT and CT groups in [2], correspondingly.

We simulate the same protocol in our model, both before and after learning. During the expected period, both prediction and stimulus inputs are active, whereas during the mismatch period, the stimulus input is paused while the prediction input remains. In our model, both nPE and pPE neurons exhibit reduced activity during the expected condition after learning (Figure 5B), consistent with previous results (Figure 3D). However, these two neuron types respond differently to the mismatch signal. pPE neurons show a negative mismatch response, indicating that they are normally excited by the stimulus. In contrast, nPE neurons show a positive mismatch response, suggesting that the stimulus inhibits their activity. At the population level, more neurons are excited than inhibited during mismatch (Figure 5C). Before learning (Figure 5D), the strongest excitatory and inhibitory mismatch responses at the single-neuron level are similar in magnitude. On average (Figure 5E), responses in the expected condition (stimulus with prediction) are comparable to those in the mismatch condition (prediction only), indicating that mismatch signals are not readily distinguishable. After learning (Figure 5F), however, the strongest excitatory mismatch responses (e.g., neuron 40) are approximately ten times larger than the strongest inhibitory responses (e.g., neuron 1). This asymmetry arises from two factors: first, responses during the expected condition are strongly suppressed; second, nPE neurons become selectively inhibited by stimulus-driven inhibition, resulting in stronger disinhibition when stimulus input is absent.

### 3.5 Mismatch response under perturbation

If the learned connectivity is optimal in computing prediction error, then any perturbations of it are suboptimal and should do worse. To test this, we probed the model by perturbing the activity of interneuron populations, mimicking a photostimulation experiment (Figure 6A).

**Fig. 6.**
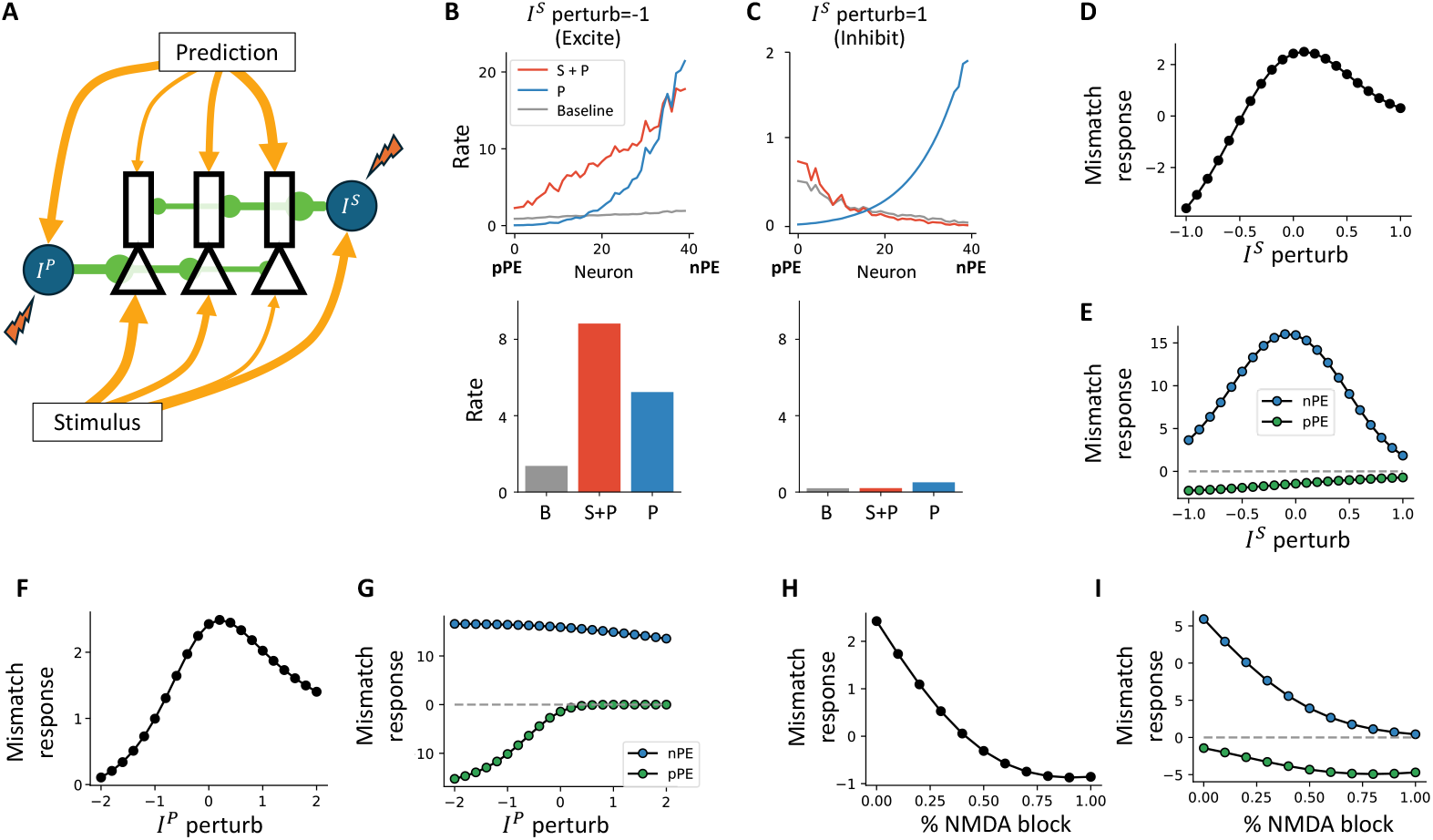
Population activity under perturbation. (A) Schematic of the task. (B) Neuronal activity when inhibiting the *I*^*S*^ population, shown at the level of individual neurons (top) and population average (bottom) across contexts. (C) Same as (B), but when exciting the *I*^*S*^ population. (D) Average mismatch response of the population across *I*^*S*^ perturbation strengths. (E) Mismatch response of a representative pPE neuron and nPE neuron across *I*^*S*^ perturbation strengths. (F, G) Same as (D, E), but for perturbing the *I*^*P*^ population. (H, I) Same as (D, E), but for blocking NMDA receptors.

By inhibiting the *I*^*S*^ population, the pyramidal neuron population becomes disinhibited (Figure 6B). However, because inhibition from *I*^*S*^ is aligned with prediction-driven excitation, nPE neurons experience stronger disinhibition than pPE neurons. This asymmetry is reflected in the increasing activity across neurons along the neuron index, which is sorted by the stimulus affinity. As a result, nPE neurons no longer compute prediction error accurately: they exhibit elevated responses in the expected condition. Consequently, the change of population response stops reflecting the prediction error. In the opposite scenario, excitation of the *I*^*S*^ population leads to widespread inhibition across the pyramidal neuron population (Figure 6C). Under this condition, nPE neurons are strongly inhibited in both the expected and mismatch cases. However, accurate mismatch computation requires nPE neurons to show increased activity in the mismatch condition, and this differential response is lost under excessive inhibition. As a result, nPE neurons fail to compute the prediction error accurately in this case as well. When plotting the mismatch response as a function of perturbation strength (Figure 6D; additional examples shown in Supplementary Figure 14A, B), we observe that performance is optimal at zero perturbation, indicating that any deviation impairs the network’s ability to compute prediction error. This degradation in performance is primarily driven by nPE neurons, rather than pPE neurons (Figure 6E), consistent with the fact that nPE neurons receive stronger inhibition from the *I*^*S*^ population.

In addition to perturbing the *I*^*S*^ population, perturbations of the *I*^*P*^ population (Figure 6F) yield qualitatively similar outcomes. However, the specific mechanisms underlying performance degradation differ across conditions. When the *I*^*P*^ population is inhibited, pPE neurons become disinhibited in the expected condition (Figure 6G, Supplementary Figure 14C), resulting in spurious responses that distort prediction error signaling. When the *I*^*P*^ population is excited, the example pPE and nPE neurons appear less affected (Figure 6G), but nPE neurons that receive both prediction and stimulus inhibition (e.g., neurons 20 to 30) become excessively inhibited and cease contributing to the population-level prediction error signal (Supplementary Figure 14D).

The impairment of predictive coding by NMDA receptor blockade is widely observed across species [35, 20, 59]. To test this in our model, we block NMDA channels on all excitatory connections. As expected, this impairs mismatch computation (Figure 6H), due to reduced excitatory drive across neurons, leading to a general decrease in response amplitude (Figure 6I, J; Supplementary Figure 14E, F). Despite the differing mechanisms, the same principle holds: any perturbation disrupts prediction error computation.

This concept of optimization can be further tested in a conjugated task, where a mismatch signal is introduced by pausing the prediction input in the model (Figure 7A). Such a mismatch could arise experimentally through optogenetic suppression of axons projecting from motor areas to the visual cortex, thereby disrupting the prediction signal.

**Fig. 7.**
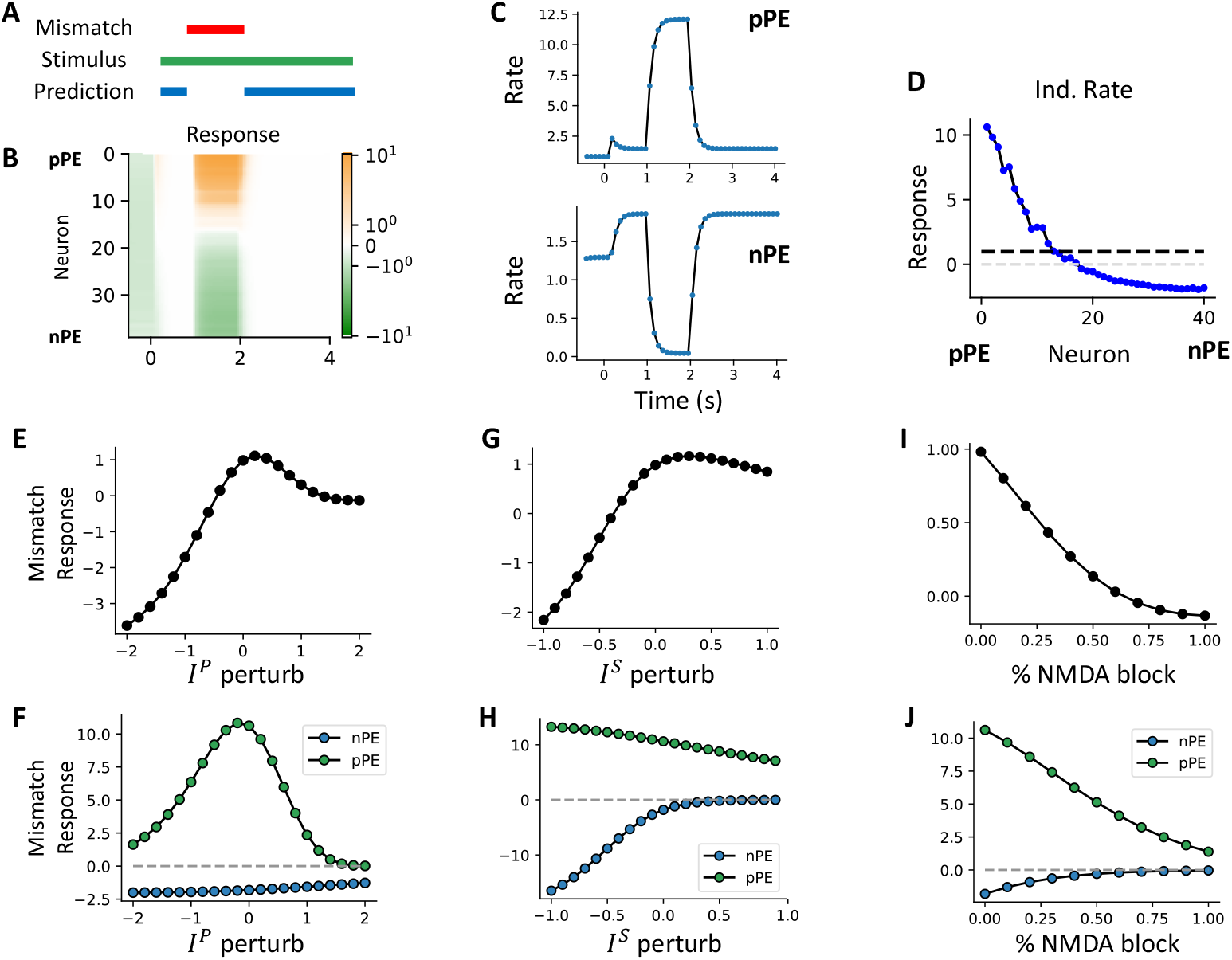
Behavior in a conjugated motor-visual mismatch task. (A) Schematic of the simulation. A mismatch is introduced by pausing the prediction signal in the model. (B) Neuronal response in the task. pPE neurons show a positive mismatch response, while nPE neurons exhibit a negative mismatch response. (C) Example neuron dynamics of a pPE and an nPE neuron. (D) Mismatch responses across individual neurons. The population average is shown as the dashed black line. (E) Population-averaged activity across *I*^*P*^ perturbation strength. (F) Mismatch response of a representative pPE and nPE neuron under *I*^*P*^ perturbation. (G, H) Same as (E, F), but for perturbing the *I*^*S*^ population. (I, J) Same as (E, F), but for blocking NMDA receptors.

In contrast to the previous scenario, pPE neurons exhibit a strong positive response to the mismatch, while nPE neurons show a modest negative response (Figure 7C). Across the neuronal population, the mismatch response gradually decreases from positive to negative values, although the average response remains positive (Figure 7D). When examining mismatch responses as a function of perturbation strength, the population response is again optimized in the unperturbed condition:

when neither the *I*^*S*^ nor the *I*^*P*^ populations, nor the NMDA channels, are manipulated (Figure 7E, G, I). The mechanisms underlying performance degradation under perturbation are similar to those described earlier (Figure 7F, H, J, Supplementary Figure 15). For example, inhibiting the *I*^*P*^ population leads to strong disinhibition of pPE neurons in the expected condition, reducing the difference between the expected and stimulus-only responses (Figure 7F, Supplementary Figure 15A). As a result, the population-level mismatch response is diminished (Supplementary Figure 15B). This mechanism is analogous to the effect of inhibiting the *I*^*S*^ population in the original mismatch task.

### 3.6 Bimodal correlation distribution emerges after learning

Our model predicts that, after learning, prediction-driven excitation should be balanced by stimulusdriven inhibition, and stimulus-driven excitation should be balanced by prediction-driven inhibition. These relationships between excitation and inhibition can be revealed by examining the correlation between neuronal responses and varying levels of stimulus and prediction input. Here, we focus on how these correlations change as a result of learning.

We first fix the prediction input strength while randomly varying the stimulus input each second (Figure 8A). In our model, before learning, a subset of neurons exhibit stronger responses when both stimulus and prediction inputs are present, compared to either input alone (Figure 3A; approximately from neuron 15 to 25). Indeed, two example neurons (Figure 8B; neuron 15 and neuron 25) show positive correlations with stimulus input. After learning, however, almost no neurons maintain elevated responses when both inputs co-occur (Figure 3C). After learning, neuron 25 switches from a positive to a negative correlation with stimulus strength (Figure 8B). We then reverse the setup by fixing the stimulus input while randomly varying the prediction input (Figure 8C). Before learning, both example neurons again show positive correlations with the prediction strength. After learning, neuron 15 switches to a negative correlation with prediction strength (Figure 8D).

**Fig. 8.**
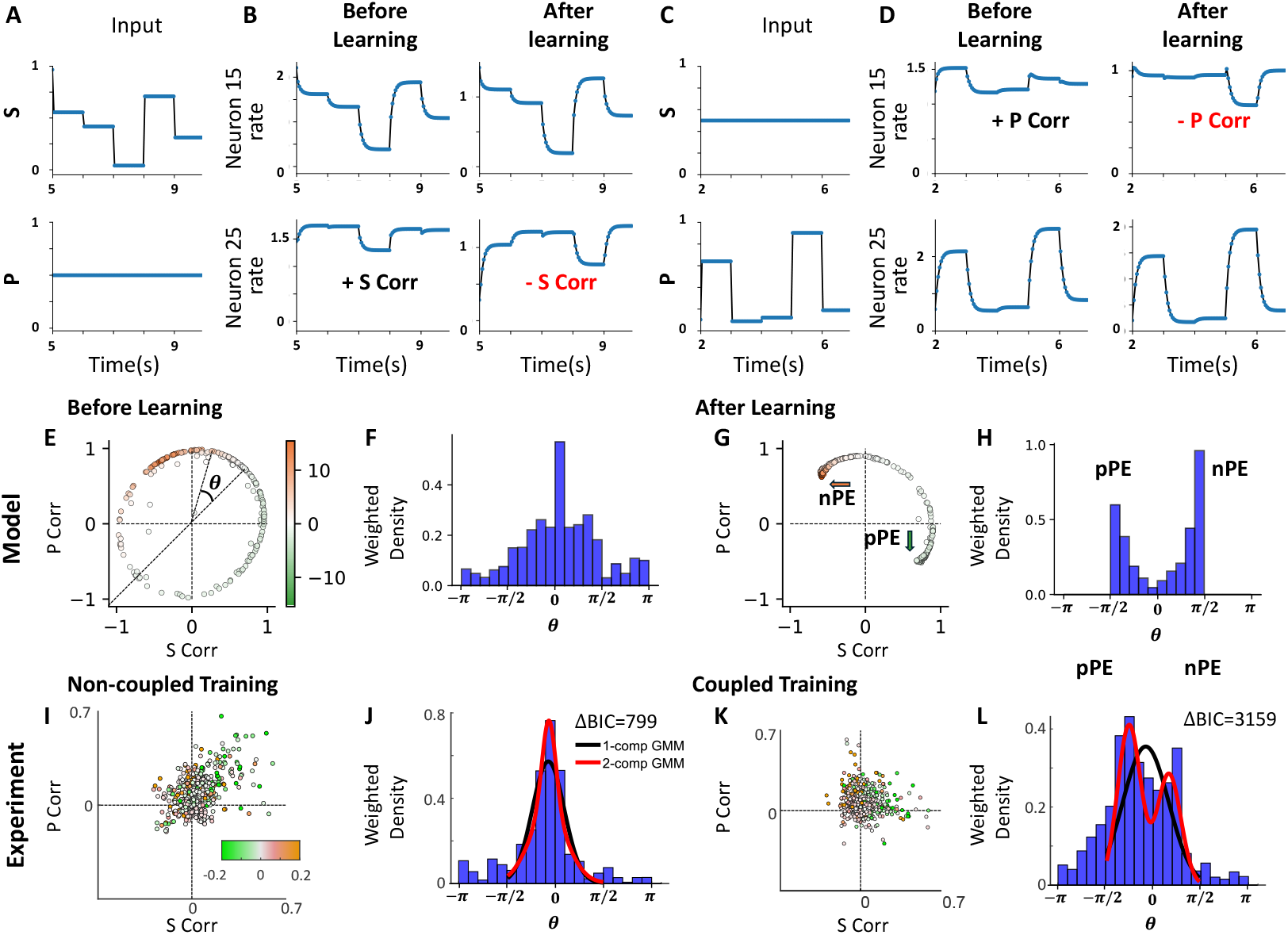
Bimodal distribution of correlation emerges after learning. (A) Stimulus input is varied across trials (top), while prediction input is fixed (bottom). (B) Responses of two example neurons (Neuron 15 and 25) to the varied stimulus input, before and after learning. The firing rate of neuron 25 changes from positively to negatively correlated with stimulus strength after learning. (C) Reversed input conditions compared to (A): stimulus input is fixed (top), while prediction input is varied (bottom). (D) Responses of the same two neurons to varied prediction input before and after learning. The firing rate of neuron 15 changes from positive to negative correlation with prediction strength. (E) Correlation of firing rates with prediction and stimulus before learning across all neurons in the model. Here, both stimulus and prediction input are varied. The color scale represents the mismatch response of individual neurons after learning. (F) Distribution of the weighted angular difference between stimulus and prediction correlations (see Methods). A value of zero indicates identical correlation with both inputs. (G, H) Same as (E, F), but after learning. A value near *π/*2 indicates positive correlation with stimulus and negative correlation with prediction. The color for a neuron is the same in (E) and (G). (I–L) Reanalysis of experimental data from [2] shows a similar bimodal distribution in the coupled training condition, but not in the non-coupled condition. The color scale represents the z-scored firing rate of individual neurons. Here, the color represents the mismatch response for that group. *N*_CT_ = 939, *N*_NT_ = 690. The red and black line in (J, L) suggests the fitted Gaussian mixture model (GMM) with one or two components (see methods and Supplementary Figure 16 for details). The difference of the Bayesian Information Criterion (ΔBIC) is used to determine which model is more favorable. ΔBIC*>* 10 suggests strong evidence that the 2-component GMM is more favorable [52]. Although fitting both CT and NT favors two-component GMMs, only the fitting of the CT dataset shows two distinct peaks in the fitted curve, indicating a true bimodal distribution.

**Fig. 9.**
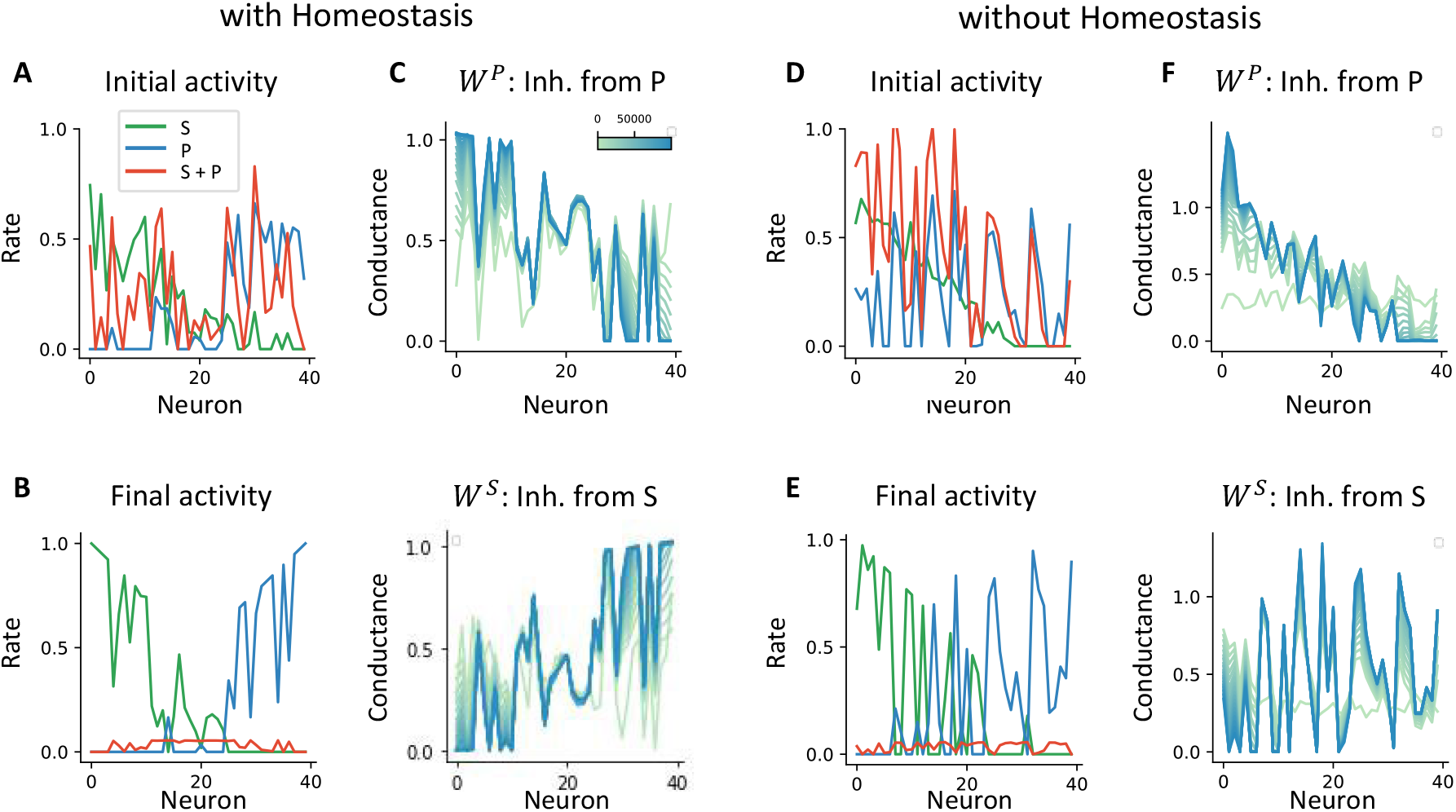
Synaptic weights and activity profiles with and without the homeostasis hypothesis, sorted by the stimulus input strength to the postsynaptic ReLU neuron. Related to Figure 2. (A–C) With the homeostasis hypothesis. (A) Initial neuronal activity in response to stimulus only (blue), prediction only (green), and both simultaneously (red). (B) Final activity after learning. (C) Inhibitory connectivity strength: *W* ^*P*^ relaying inhibition from prediction (top), and *W* ^*S*^ relaying inhibition from stimulus (bottom). Different colors represent weights at different training stages; a darker color indicates a later stage of learning. (D–F) Without the homeostasis hypothesis. Same format as (A–C), but the simulation is performed by shuffling the prediction input, removing the anti-correlation between prediction and stimulus input strength.

**Fig. 10.**
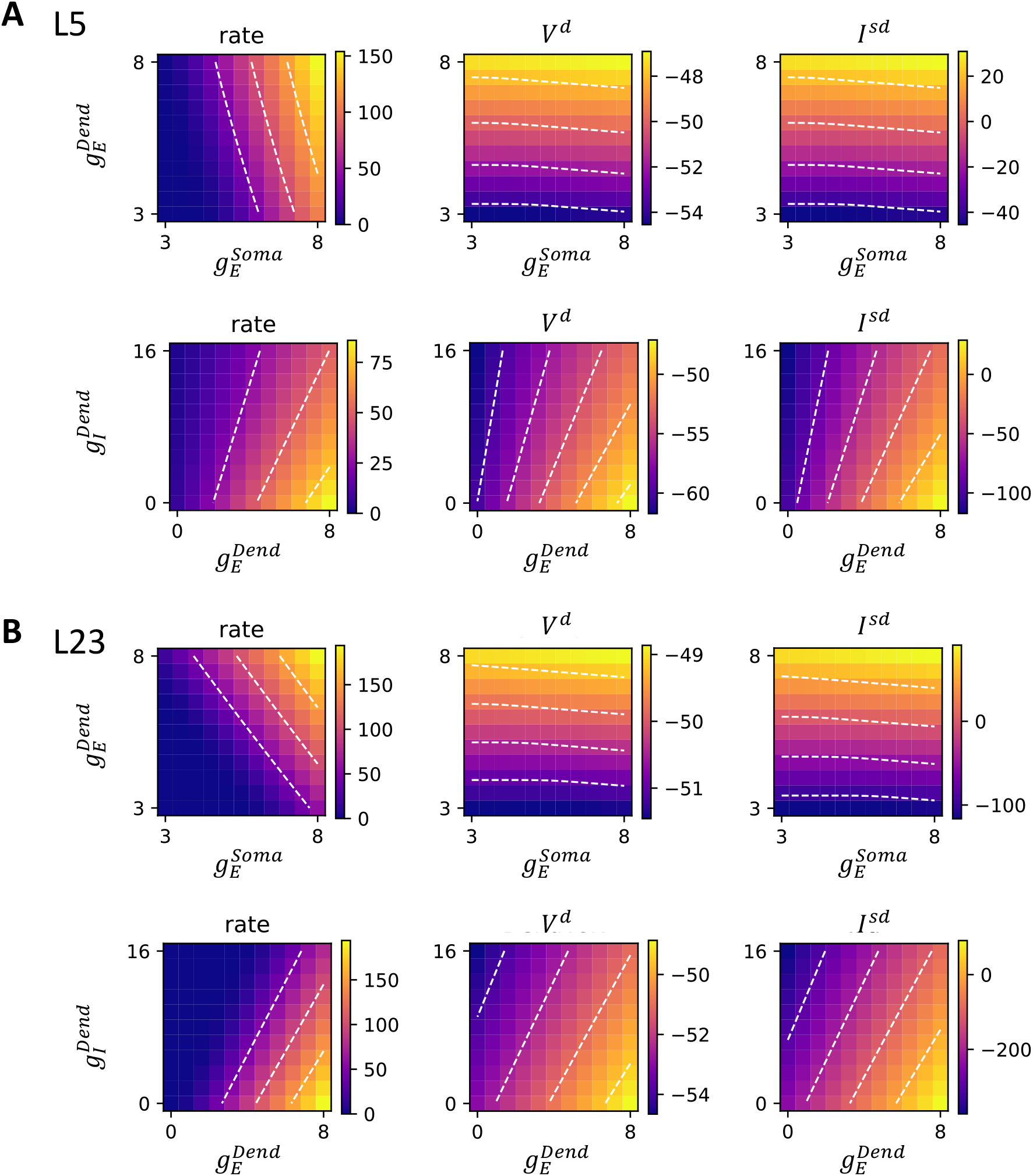
Activity of the two-compartment hybrid pyramidal cell model. Related to Figure 3. (A) Layer 5 pyramidal cell model. The first row shows firing rate, dendritic voltage, and coupling current as functions of somatic excitation 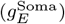 and dendritic excitation 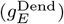). White dashed contours indicate iso-response levels. Dendritic input has a strong influence on dendritic voltage and coupling current, but not on somatic firing rate. The second row illustrates the interaction between dendritic excitation 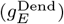 and dendritic inhibition 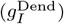. The slope of the contour lines is steepest when inhibition dominates, suggesting a divisive effect; when excitation dominates, inhibition acts more subtractively. (B) Same analysis for a layer 2/3 pyramidal cell model. In this model, coupling strength is much higher than in the layer 5 model, resulting in dominantly subtractive inhibition in this parameter regime.

**Fig. 11.**
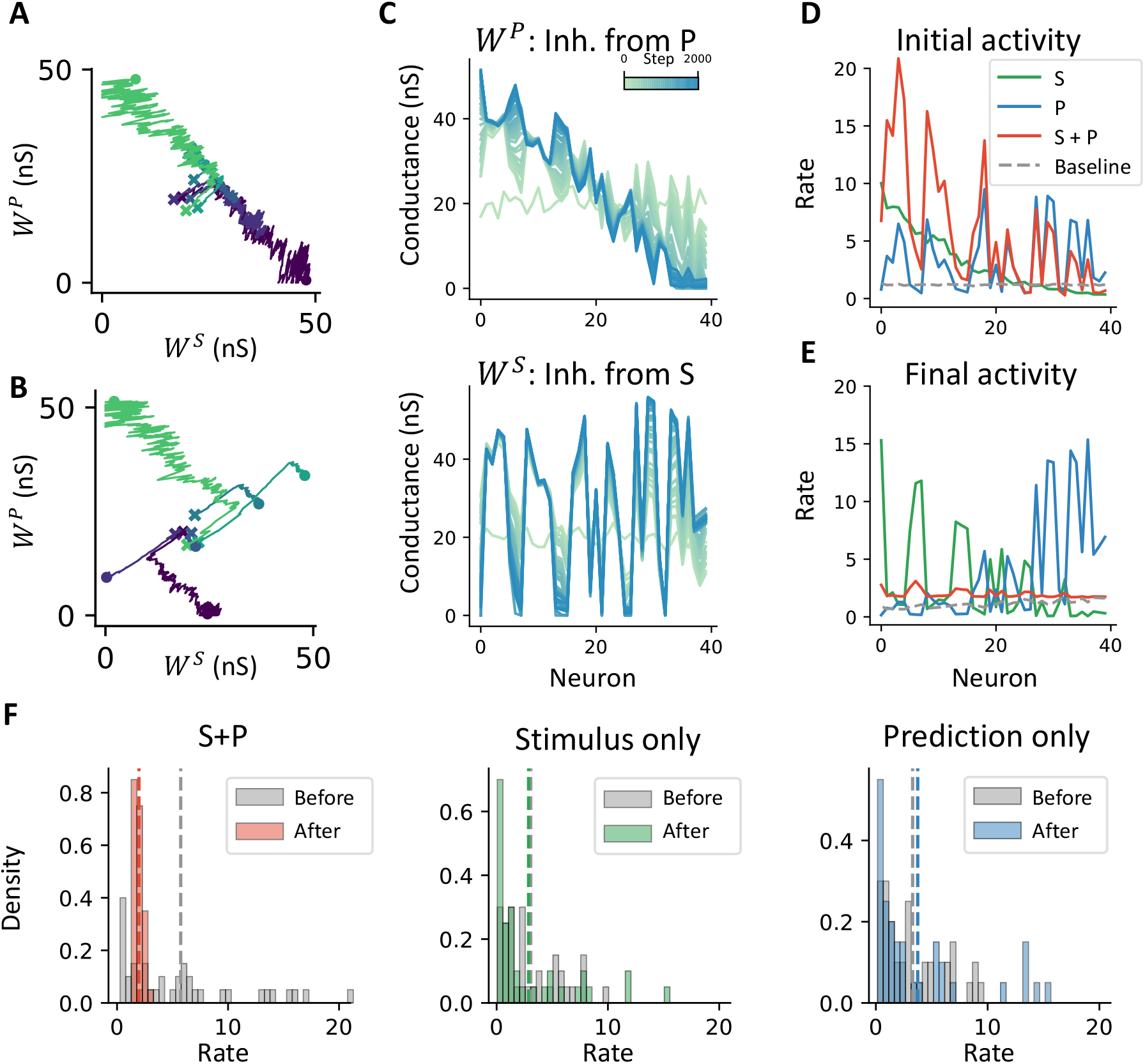
Learning does not require the homeostasis hypothesis in the realistic neuron model. Related to Figures 3. (A) Weight updates in the realistic neuron model under the homeostasis hypothesis. (B) Same as (A), but without the homeostasis hypothesis. (C) Final inhibitory connectivity strength from *I*^*P*^ (top) and *I*^*S*^ (bottom). (D) Initial neuronal activity crosses different contexts before learning. (E) Final neuronal activity after learning. (F) Activity distribution before and after learning. Population activity decreases only when both prediction and stimulus are presented. The results qualitatively match those shown in Figure 3.

**Fig. 12.**
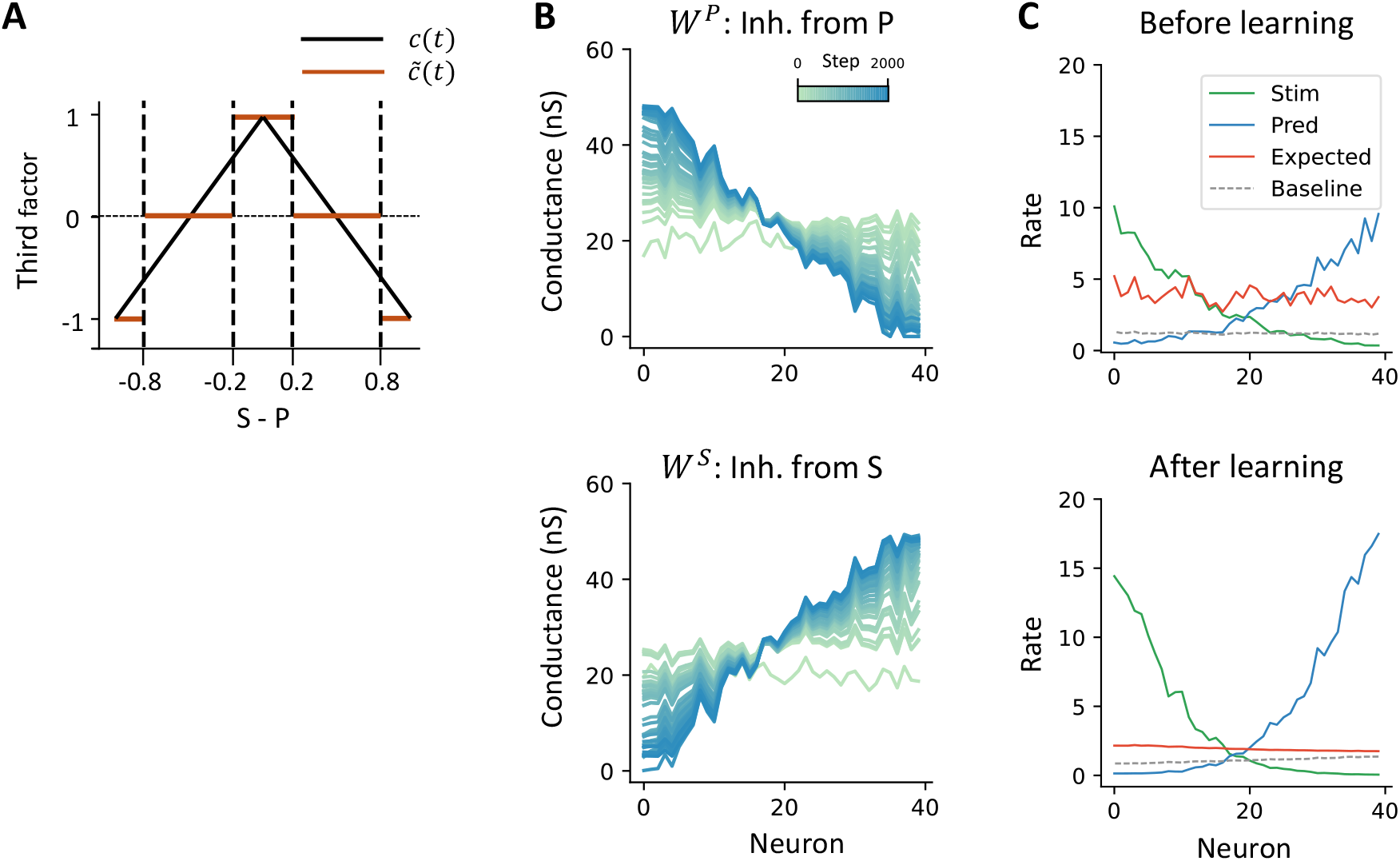
Estimated third factor achieves the same qualitative connectivity. Related to Figures 3. (A) The original third factor *c*(*t*) and estimated piece-wise third factor 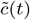 (B) Learned Connectivity weight *W* ^*P*^ and *W* ^*S*^, using the estimated third factor 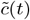. (C) Neuron activity before and after learning.

**Fig. 13.**
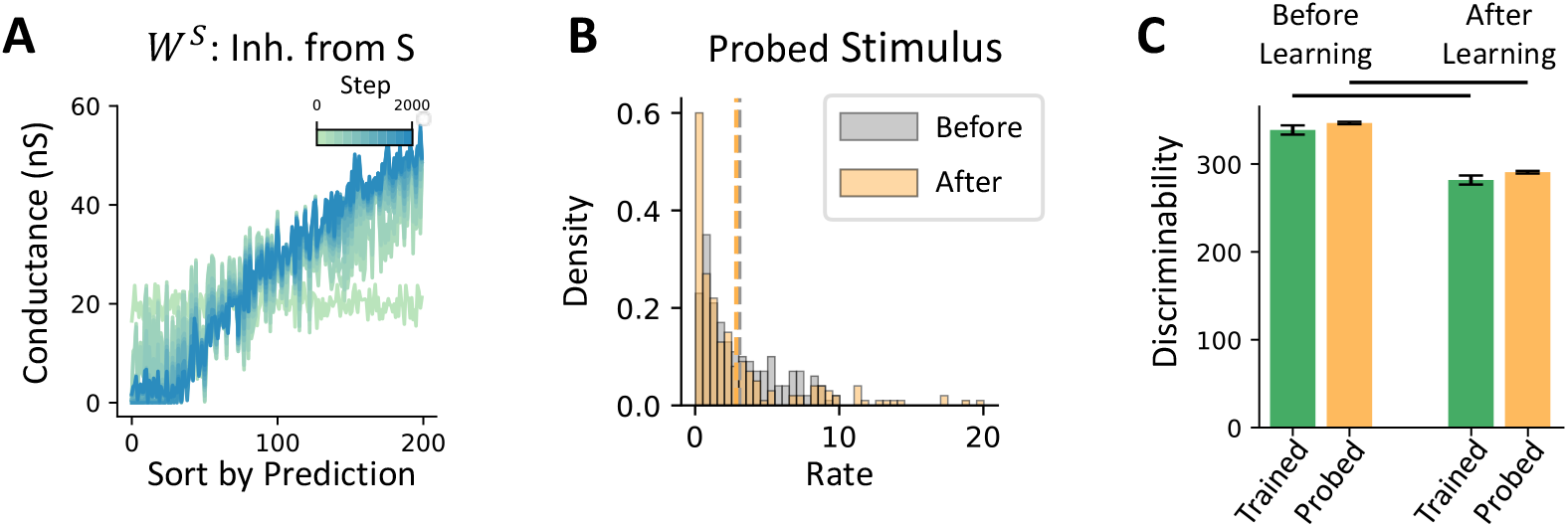
Analysis related to the probed stimulus. Related to Figure 4. (A) Inhibitory connectivity from the stimulus (*W* ^*S*^) is aligned with the prediction input. (B) Activity distribution in response to the probed stimulus before and after learning. (C) Discriminability between the trained and probed stimuli, measured before and after learning in the absence of prediction input. Discriminability is quantified as the sum of squared d-prime values across all neurons (see Methods), averaged across another twenty probed stimuli. Each probed stimulus is generated by shuffling the trained stimulus across neurons. No significant difference in discriminability is observed between the trained and probed stimuli (*p >* 0.1). Learning reduces discriminability between stimulus pairs by approximately 10% (*p <* 0.001), suggesting a trade-off between predictive error computation and stimulus-specific decoding ability.

**Fig. 14.**
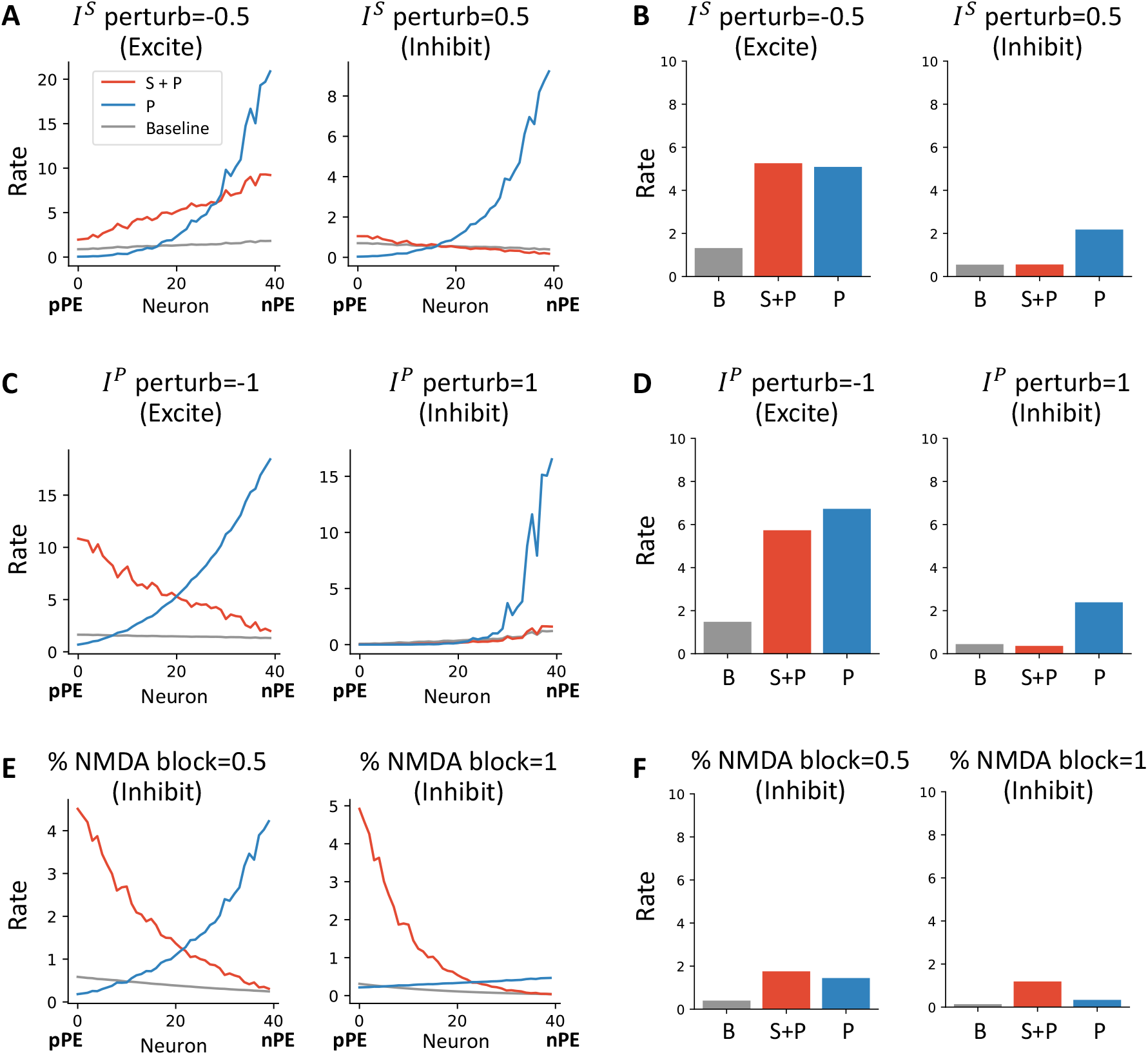
Examples of perturbation. Related to Figure 6. (A) Firing rates of individual neurons when perturbing *I*^*S*^ with strengths *I*^*S*^ perturb = −0.5 and 0.5. The bracket “excite” and “inhibit” suggest the effect on disynaptic pyramidal cells. (B) Corresponding population-averaged firing rates. (C, D) Same as (A, B), but for perturbing *I*^*P*^ with strengths *I*^*P*^ perturb = 1 and 1. (E, F) Same as (A, B), but for blocking NMDA receptors at proportions 0.5 and 1.

**Fig. 15.**
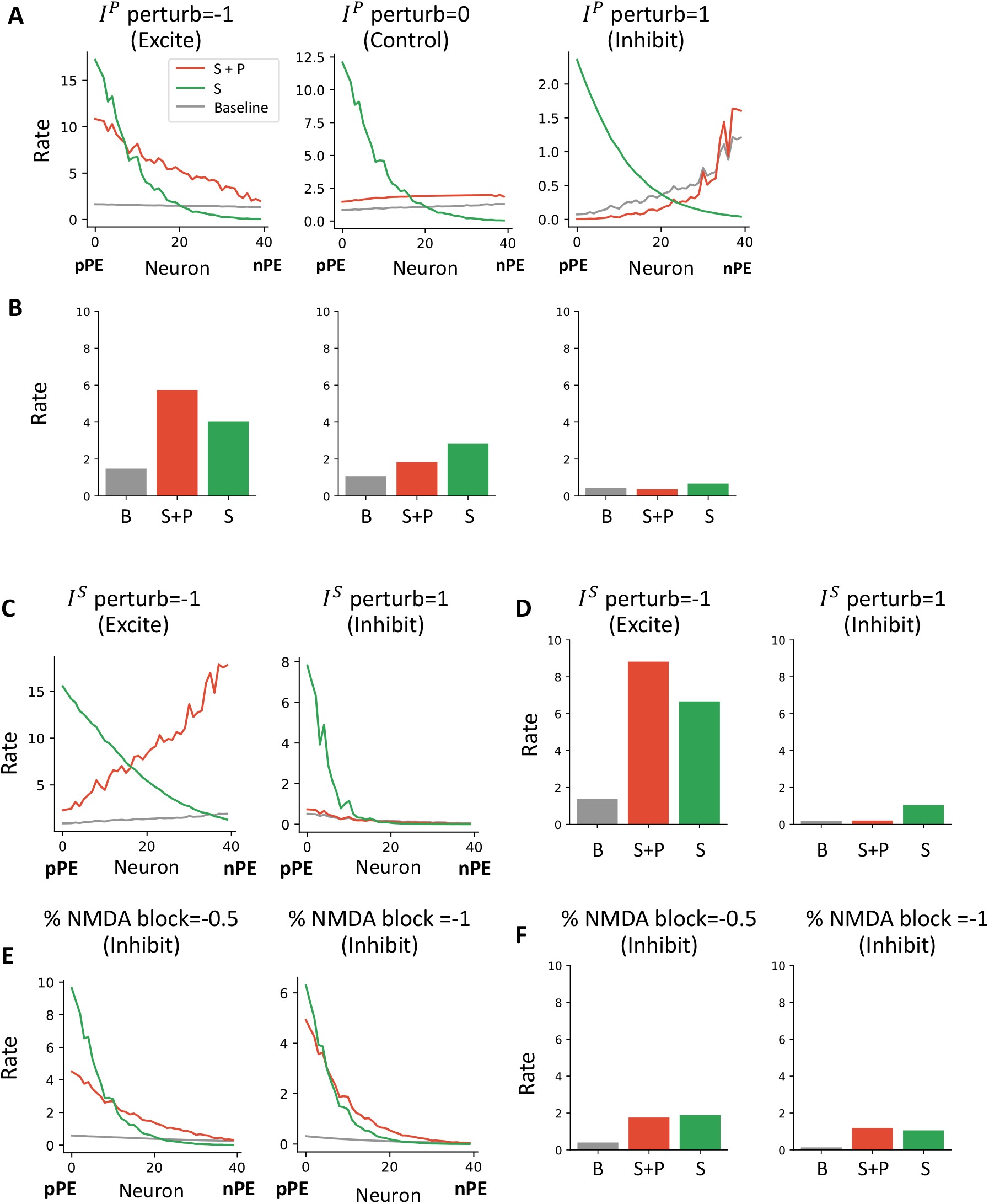
Perturbed activity in the conjugated motor-visual mismatch task. Related to Figure 7. (A) Activity of individual neurons when perturbing the *I*^*P*^ population with strengths *I*^*P*^ perturb = −0.5, 0 (control), and 0.5. (B) Corresponding population-averaged activity. (C, D) Same as (A, B), but for perturbing the *I*^*S*^ population with strengths *I*^*S*^ perturb = −1 and 1. (E, F) Same as (A, B), but for blocking NMDA receptors at proportions 0.5 and 1.

We next vary the stimulus and prediction input strengths simultaneously and compute each neuron’s response correlation with stimulus and prediction input separately. To improve statistical reliability, we use a larger model without the homeostasis constraint, with *N* = 200 neurons and a continuous simulation duration of 200 seconds. In Figure 8E, each dot represents a single neuron, with its correlation with stimulus input on the x-axis and correlation with prediction input on the y-axis. The color indicates the mismatch response after learning. The dots lie approximately on a unit circle, suggesting that the variance in neural responses can be fully explained by the combination of stimulus and prediction input.

We define the angular deviation *θ* for each neuron as the difference between the angle of its correlation vector in polar coordinates and *π/*4. A value of *θ* = 0 indicates equal correlation with stimulus and prediction input. We further weight each *θ* by the radius of the corresponding point in polar coordinates and plot the weighted distribution of *θ*. Before learning, *θ* follows a unimodal distribution centered near zero (Figure 8F). After learning, however, the correlation points cluster in the second and fourth quadrants, consistent with a functional split between nPE and pPE populations (Figure 8G). This separation is further reflected in the emergence of a bimodal distribution in the weighted distribution of *θ* (Figure 8H).

Finally, we test whether the same bimodal correlation structure emerges in experimental data. We analyze previously published recordings from [2] focusing on both non-coupled training (NT) and coupled training (CT) mice during the open-loop condition, in which movement and visual flow are not coupled. As in the model, we compute the correlation between neuronal responses and either visual flow (stimulus) or locomotion speed (prediction), and represent these in polar coordinates (Figure 8I, K). We then plot the weighted distribution of the angular difference *θ* as defined earlier (Figure 8J, L). In the NT group, the distribution of *θ* is unimodal and centered near zero, indicating similar correlation with stimulus and prediction inputs. In contrast, the CT group shows a bimodal distribution, suggesting that the functional separation into pPE and nPE neurons emerges only with coupled sensorimotor experience, consistent with our model predictions. This bimodality is further confirmed by fitting the angular data to Gaussian mixture models (GMM) with one or two components (see Methods; Figure 8J, L, Supplementary Figure 16).

**Fig. 16.**
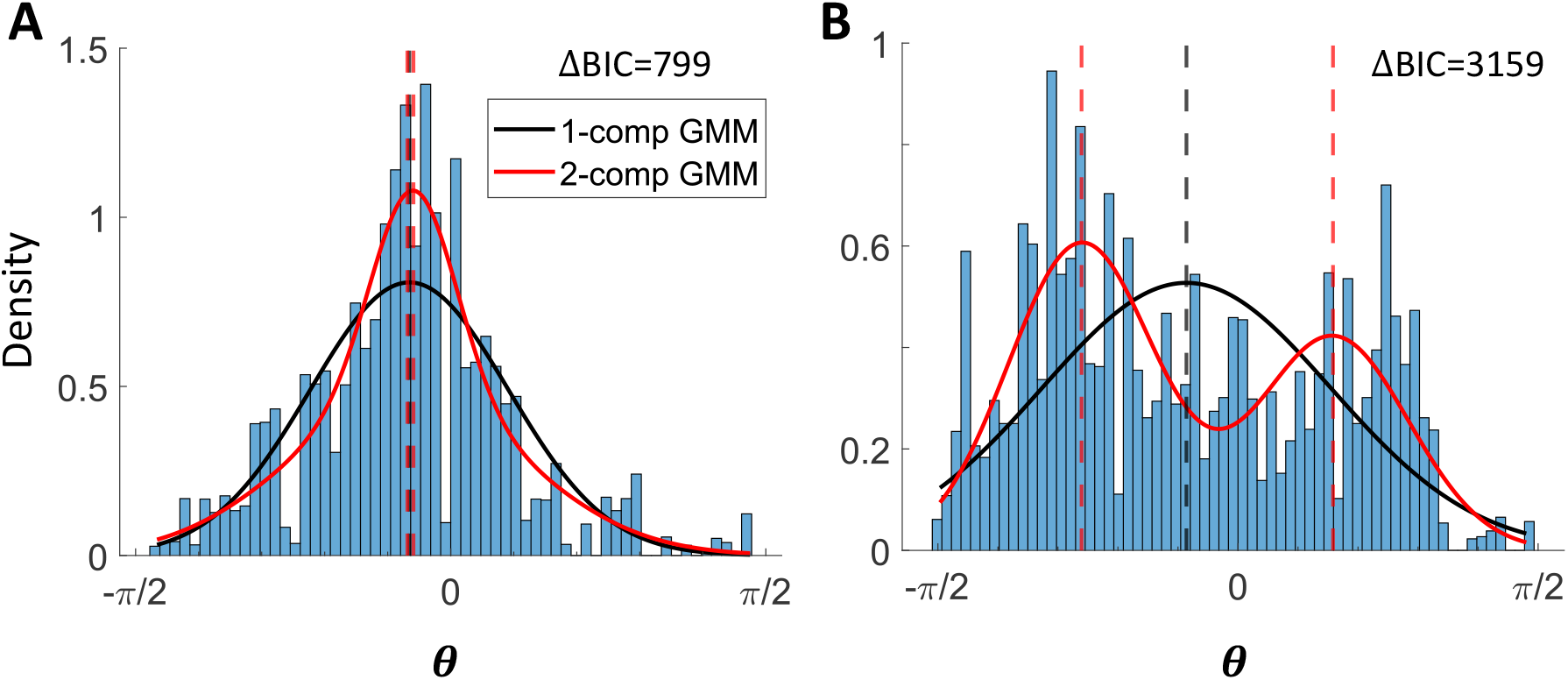
Fitting pseudo-data to Gaussian Mixture Model with one or two components (GMM). Related to Figure 8. See Methods. The difference in the Bayesian Information Criterion (BIC) suggests which model is more favorable. The fitted lines are identical to those in Figure 8. (A) GMM fit to the pseudo data generated from the non-coupled training data. Solid lines represent the fitted distribution; dashed lines indicate the means of the Gaussian components. (B) GMM fit to the pseudo data generated from the coupled training data.

## 4 Discussion

Our work proposes that a three-factor learning rule optimizes local inhibitory connectivity for the computation of prediction errors in sensory areas. Each component of the rule is biologically supported, essential for the learning process, and converges to the same optimal solution as the gradient descent algorithm. After learning, positive prediction error (pPE) and negative prediction error (nPE) neurons emerge simultaneously, and their population activity reflects the prediction error: the difference between prediction and stimulus input. Because error is minimized only for stimuli that have been paired with a prediction signal, the model captures the selective modulation of responses observed in auditory cortex [4, 3]. We reproduce the response in a motor-visual mismatch task [2] and further argue that any perturbation to the optimized circuit impairs performance. Finally, the model predicts a bimodal distribution of correlation with stimulus and prediction inputs after learning, a pattern that is confirmed by reanalysis of existing data.

The three-factor learning rule, also known as the neoHebbian learning rule, was first introduced by Gerstner and colleagues [21, 24, 37]. It extends traditional Hebbian plasticity [27] by incorporating a third factor in addition to pre- and postsynaptic activity. This third factor can represent signals such as reward, punishment, or surprise, and is often mediated by neuromodulators like dopamine [10, 45, 8, 53]. In the three-factor framework, the pre- and postsynaptic activity sets a “tag” at the synapse, marking it as eligible for modification. The third factor can then arrive later (within a certain time window) to modify the actual synaptic weight using that tag. In doing so, three-factor learning bridges the timescales between fast neuronal activity (milliseconds) and slower behavioral outcomes (seconds) [24, 64]. Moreover, any biologically realistic learning mechanism must address the stability-plasticity dilemma [41], in which high plasticity leads to forgetting and high stability impairs new learning. The third factor has been proposed as a solution to this dilemma [26, 6], by dynamically gating plasticity in a context-dependent manner.

In the context of motor–sensory mismatch, animals are typically trained in a passive setting to associate a specific motor output with a particular stimulus. This approach minimizes confounds from other top-down signals such as attention [2, 4]. Only recently has an unsigned prediction error been observed in the activity of the locus coeruleus axons, presumably reflecting noradrenaline release in sensory areas [32, 30]. Based on this, we propose that noradrenaline acts as the third factor to optimize local inhibitory connectivity for prediction error computation in sensory circuits. Also, serotonin may play a similar modulatory role in guiding synaptic plasticity and local circuit optimization [63].

In our model, the three-factor learning rule is applied at local inhibitory synapses onto pyramidal cells, but not at the excitatory synapses onto interneurons. We omit the latter for three reasons. First, mismatch responses in interneurons do not appear to depend on experience, as reported in [2]. Second, successful balancing of excitation via relayed inhibition requires that interneurons be stimulus-selective, a condition that may not be biologically feasible given their dense and non-selective connectivity [33] (but see [65]). Third, optimizing excitatory synapses onto interneurons would require access to the firing rates of their downstream pyramidal targets, implying a non-local learning rule. Nevertheless, the potential role of plasticity of excitatory synapses onto interneurons may contribute to circuit flexibility in ways not addressed here.

By leveraging the framework of the three-factor learning rule, our model simplifies the assumptions required for the emergence of nPE and pPE neurons, relative to previous work [28]. To our knowledge, that study is the first to propose a mechanism by which nPE and pPE neurons emerge simultaneously within the same circuit. In their model, co-emergence relies on establishing excitation–inhibition balance at both dendritic and somatic compartments through inhibitory plasticity. However, achieving this outcome depends on several specific assumptions. First, the model requires two distinct PV interneuron subtypes, each selectively receiving either stimulus or prediction input. Second, the learning scheme is not entirely local, raising questions about its biological feasibility. Third, dendritic input can only excite the connected soma but cannot exert inhibition. In contrast, we show that a biologically plausible, fully local three-factor learning rule is sufficient to reproduce the emergence of nPE and pPE neurons without these cumbersome assumptions.

Our simplification can be understood as a shift in the objective of circuit optimization. The studies by [28, 29] propose that neural circuits aim to minimize overall firing rates to conserve energy, consistent with the original formulation of predictive coding [47]. In their framework [28], the error signal arises as a by-product of the circuit’s limited experience with mismatched inputs, rather than as an explicitly learned feature. In contrast, we argue that the neuronal population is explicitly optimized to compute the prediction error with the correct amplitude in mismatched conditions, rather than merely suppressing activity in expected cases. Indeed, we demonstrate mathematically (see Supplementary Materials) that the optimal local connectivity can only be achieved through learning driven by error in mismatched conditions.

However, our model assumes that the animal can estimate a global error signal in some form, which could raise concerns about circular reasoning: if prediction error drives learning, how can it be computed before an accurate internal model exists? Yet, studies show that passive exposure is sufficient for animals to generate prediction error-like signals [2, 4]. This suggests that animals may rely on approximate surprise signals, such as arousal responses to unexpected events [32], rather than precise prediction error estimates. Theoretically, this arousal signal could be derived by tracking variations in sensory input over longer timescales, such as across training sessions, and may serve as a latent predictive signal that guides the learning process [26]. Though such arousal signals may not represent the true global error, we show that an approximation is sufficient to guide local synaptic optimization (Supplementary Figure 12). Still, the mechanisms by which animals generate these signals remain an important topic for future investigation.

Functionally, we argue that learning in sensory areas serves to reduce cognitive load, the mental resources required to perform a task, during prediction error computation [57, 58]. For example, when first learning to ride a bicycle, one must consciously monitor every movement to avoid falling. With experience, however, bicycling becomes effortless, even on uneven terrain. Over time, the skill becomes automatic. Similarly, if visual and motor systems learn to compute prediction error locally, as our model proposes, this may offload computation from higher-order areas, such as the prefrontal cortex, and thereby conserve cognitive and metabolic resources. Supporting this view, an fMRI study of humans learning a visual–motion association task shows reduced activity in prefrontal regions in the later stages of learning compared to the early stages [5]. Future work should investigate whether similar signatures emerge in mice trained on predictive coding tasks.

To dissect the circuitry underlying predictive coding, it is essential to selectively target and perturb each component of the model. Recent studies suggest that Layer 2/3 *Rrad* and *Baz1a* pyramidal neurons are enriched for pPE responses, whereas *Adamts2* neurons preferentially exhibit nPE responses [43, 12]. These transcriptomic subtypes may also possess distinct electrophysiological properties [31]. Consistent with this, we demonstrate mathematically that different initial conditions, defined by gene expression, lead to distinct pPE and nPE outcomes after learning (see Supplementary Materials), providing a potential mechanism linking transcriptomic identity to learned functional roles. Local interneurons are essential for converting long-range excitation into local inhibition. Among them, somatostatin-positive (SST) interneurons are the most likely mediators of stimulus-driven inhibition, as supported by studies of motion–visual mismatch responses [2, 61, 34].

However, the identity of the interneurons that relay prediction-driven inhibition remains unclear. In the auditory cortex, parvalbumin-positive (PV) interneurons have been implicated in mediating motion-related inhibition [50, 51]. However, findings from [4] suggest that PV cells may instead implement a more uniform gain control mechanism, rather than conveying specific predictive signals. In contrast, SST interneurons have also been shown to mediate stimulus-selective inhibition [51], raising the possibility that they may also relay prediction-related inhibition. If so, the model would require functionally distinct subpopulations of SST cells to carry inhibition from different sources. Indeed, heterogeneous responses within the SST population (see Fig. S3 in [2]) support this possibility. Another major interneuron subtype, vasoactive intestinal peptide-positive (VIP) cells, primarily inhibit SST interneurons and thereby disinhibit pyramidal neurons [46], making it less likely a candidate for mediating prediction-driven inhibition. However, recent findings from our group identify a VIP subpopulation that co-expresses the neuropeptide cholecystokinin (CCK) and targets pyramidal neurons directly, with comparable synaptic strength to that onto SST cells, and receives input directly from the motor cortex [17]. These VIP/CCK cells may serve as a candidate population for mediating prediction-driven inhibition.

To elucidate the functional role of a specific interneuron subtype, our model predicts distinct signature behaviors of *I*^*S*^ or *I*^*P*^ populations under targeted perturbations. These signatures can help determine the functional identity of a given interneuron subtype. Importantly, they are not detectable at the population level, since any perturbation suppresses the overall mismatch response (Figure 6, Figure 7), but must be identified at the single-cell level. In the motion–visual mismatch task, perturbing *I*^*S*^ is predicted to reduce the response of nPE neurons (Figure 6E). In contrast, excitation of the *I*^*P*^ population increases the response of pPE neurons in the expected condition, thereby reducing the mismatch signal at the population level (Figure 6G). Additionally, inhibition of the *I*^*P*^ population leads to a functional shrinkage of the nPE population (Supplementary Figure 14C), though not necessarily a decrease in individual response amplitude. Similar patterns are expected in the conjugated task as well (Figure 7, Supplementary Figure 15). Future experiments can test these predictions using highresolution genetic tools to target specific interneuron subtypes, combined with continuous rather than binary perturbation strategies.

In general, our theory provides a unifying framework for any instance in which functionally relevant signals are compared across cortical areas, consistent with [34]. Indeed, similar nPE and pPE response patterns have been observed in the visual cortex during audio–visual mismatch [23] and in the auditory cortex during motor–audio mismatch [54]. Notably, in the audio–visual paradigm, the auditory input, serving as the predictive signal, only suppresses visual responses after learning [23], further supporting the idea that prediction-driven inhibition becomes aligned with stimulus-driven excitation through plasticity. Moreover, the functional segregation of suppressive and enhanced neural responses in the auditory cortex of vocalizing marmosets [18] can also be accounted for by the emergence of pPE and nPE neurons in our model. It is worth noting that self-generated sounds can change over time, requiring continuous plasticity to maintain accurate cancellation over age. These observations suggest that our model offers a unified framework for understanding one aspect of corollary discharge [56, 15], wherein motor-related signals are sent to sensory areas to suppress predictable reactions.

## Methods

### 4.1 Simplified Neural Circuit

We begin with a simplified neural circuit with ReLU neurons to illustrate the mechanism of our three-factor learning rule. Here, all the variables are dimensionless and are updated instantaneously without temporal dynamics.

For the *i*-th ReLU unit, activity is given by:

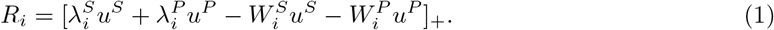

For simplicity, we omit the subindex *i* in the following. Within the above equation,*λ*^*S*^ and *λ*^*P*^ denote the excitatory input strengths from stimulus and prediction, respectively. *W*^*S*^ and *W*^*P*^ represent the inhibitory synaptic weights from interneuron populations *I*^*S*^ and *I*^*P*^, whose activity is the same as the stimulus and prediction input *u*_*S*_ and *u*_*P*_, or simply *S* and *P* when unambiguous. The operator [*x*]_+_ denotes a rectifier (ReLU) function, where [*x*]_+_ = max(0, *x*). The synaptic weights *W*^*S*^ and *W*^*P*^ are plastic and updated according to our three-factor learning rule.

In the case of homeostasis, *λ*^*S*^ + *λ*^*P*^ = 1. Thus, for *i*-th neuron, we can write 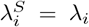 and 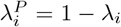. In the case of two neuron *N*_*neu*_ = 2, we set *λ*_1_ = 1, *λ*_2_ = 0. In the case of forty neuron *N*_*neu*_ = 40, the values of *λ*_*i*_ are linearly spaced from 1 to 0 across the population.

In the case without homeostasis, we shuffled the 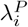 across neurons such that the total input 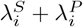 is no longer constrained to equal 1.

### 4.2 Three-factor learning rule

Our learning mechanism can optimize the mismatch calculation, as proven in the Supplementary Material.

The inhibitory connections are initiated with 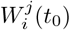 for both interneuron populations *j* = *S, P* with some jitter, mimicking the average synaptic strength from a population that should be similar across post-synaptic neurons. Again, we omit the subscript *i* in the following derivations.

These inhibitory synapses are updated with a three-factor learning rule as follows:

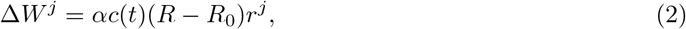

where *j* = *S, P*. Here, *R* denotes the activity of the postsynaptic pyramidal neuron, and *r*^*j*^ represents the steady-state activity of the presynaptic interneuron. *R*_0_ is a small target firing rate shared by all neurons, and *α* is the learning rate.

The third factor, *c*(*t*), is defined as:

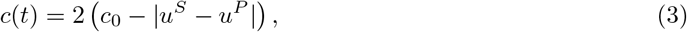

such that when *u*^*S*^ = *u*^*P*^, *c*(*t*) = 1, and when |*u*^*S*^ − *u*^*P*^ | = 1, *c*(*t*) = −1. The threshold *c*_0_ of 0.5 is chosen for simplicity. Also, our choice of *c*_0_ implies that a rough estimation of the prediction error is sufficient to serve as the third factor, since any input with *c*(*t*) *>* 0 is considered as an expected case.

We further show that an estimated piece-wise third factor is sufficient to guide the local learning. In this case,

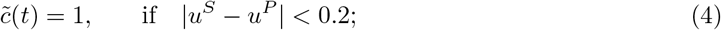

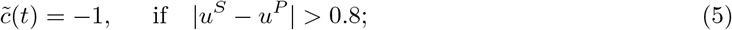

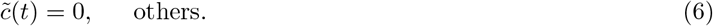

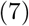

Here, the subindex *b* is short for binary. A diagram is in Figure 2D.

During training, *u*^*S*^ −*u*^*P*^ is drawn from a truncated normal distribution (Figure 2D) with mean zero and standard deviation *σ*_error_ = 0.5, reflecting the assumption that training samples are dominated by expected cases where *u*^*S*^ *≈ u*^*P*^.

The mismatch modulation saturates when the mismatch signal becomes sufficiently large, and is defined as:

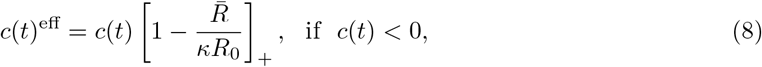

where *κ* is a scaling factor that sets the saturation level of the mismatch utility, and 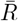 is the average activity of the neuron population.

In the case of realistic neuron models, synaptic weight updates are further capped at high conductance values:

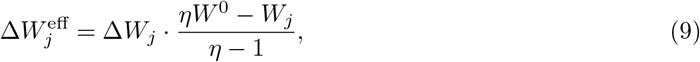

where *η >* 1 is the saturation factor for synaptic weights, and *W* ^0^ is the initial averaged synaptic strength.

### 4.3 Pyramidal neuron and interneuron models

In this work, we use a simplified hybrid dendritic model. For each pyramidal neuron, the dendritic membrane potential *V* ^*d*^ and somatic firing rate *R*^*s*^ are updated at each time step as follows:

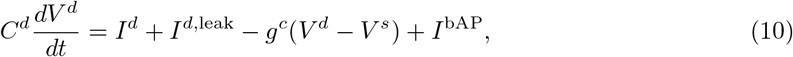

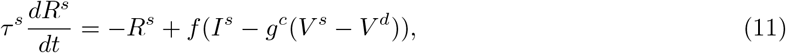

where *I*^*s*^ and *I*^*d*^ are the total synaptic currents into the soma and dendrite compartments, respectively. The dendritic leak current is given by *I*^*d*,leak^ = −*g*^*d*,leak^(*V* ^*d*^ − *V* ^*d*,rest^), where *V* ^*d*,rest^ is the resting dendritic potential and *g*^*d*,leak^ is the leak conductance. The soma and dendrite are coupled via conductance *g*^*c*^, resulting in a coupling current *I*^*sd*^ = *g*^*c*^(*V* ^*d*^ − *V* ^*s*^).

The dendritic compartment also receives input from backpropagating action potentials (bAPs), which we model as a rate-averaged current following [39]: *I*^bAP^ = −*g*^*c*^(*V* ^*d*^ − *V* ^bAP^)*t*^bAP^*R*^*s*^, where *V* ^bAP^ is the averaged membrane potential during a spike, *t*^bAP^ is the duration of the bAP, and *R*^*s*^ is the firing rate at the soma. We assume that somatic membrane potential fluctuations are negligible between action potentials, and therefore take *V* ^*s*^ *≡ V* ^*s*0^ to be constant.

The activation function *f* that converts somatic input current to firing rate is taken from [1]:

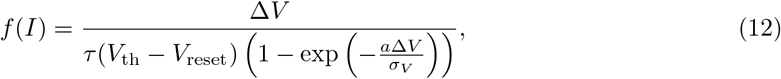

where Δ*V* = *I/g*_*L*_ + *V*_*l*_ − *V*_*th*_.

The model parameters are selected based on prior experimental work [44, 39]. The coupling conductance *g*_*c*_ is set to a small value for neurons with extensive dendritic trees, such as Layer 5 pyramidal cells, where our single-neuron model reproduces shunting dendritic inhibition (Supplementary Figure 10A). In contrast, *g*_*c*_ is set higher for neurons with more compact dendritic arbors, such as Layer 2/3 pyramidal neurons, where dendrites are closer to the soma and dendritic inhibition exhibits primarily subtractive effects (Supplementary Figure 10B).

We use a point-neuron model to describe the average activity of each interneuron population:

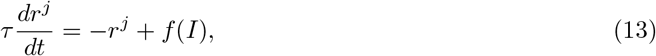

where *j* = *S, P* denotes the stimulus- or prediction-driven interneuron population, and *f* (*I*) is the same activation function defined previously.

### 4.4 Synapse model

We use a conductance-based synaptic model in our simulations, following [62]. The AMPA and GABA synapses are described by:

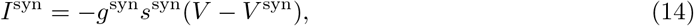

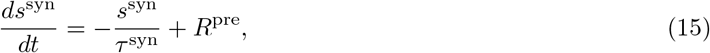

where syn = *A, G* denotes AMPA and GABA synapses, respectively. Here, *g*^syn^ is the maximal synaptic conductance, *s*^syn^ is the gating variable representing the fraction of open channels, *V* ^syn^ is the synaptic reversal potential (determined by whether the synapse is excitatory or inhibitory), *τ* ^syn^ is the synaptic time constant, and *R*^pre^ is the presynaptic firing rate.

The NMDA synapse model includes both a voltage-dependent magnesium block, *f* ^Mg^(*V*), and a saturating gating variable, *s*^*N*^ :

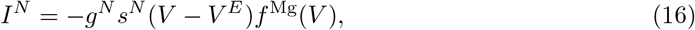

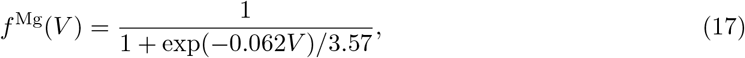

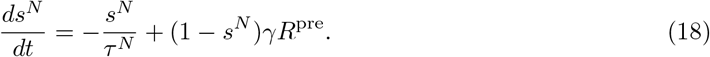

Here, *g*^*N*^ is the maximal NMDA conductance, *V* ^*E*^ is the excitatory reversal potential, *τ* ^*N*^ is the NMDA time constant, *γ* is a saturation scaling factor, and *R*^pre^ is the presynaptic firing rate. The function *f* ^Mg^(*V*) captures the voltage-dependent magnesium block characteristic of NMDA receptors.

The stimulus input to both pyramidal neurons and the first interneuron population *I*^*S*^ is modeled as an excitatory conductance to the soma, denoted by 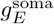. When illustrating the behavior of individual pyramidal neurons in Supplementary Figure 10, the dendritic compartment additionally receives excitatory and inhibitory input conductances, denoted by 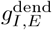. Because our model employs conductance-based synapses with distinct reversal potentials for excitation and inhibition, the total input current is not simply a linear difference between excitatory and inhibitory conductances. This formulation allows us to capture more complex and dynamic behaviors in the circuit.

### 4.5 Input and network connectivity

We include only the necessary connections required for the model. A schematic of the network architecture is shown in Figure 1E. The maximum synaptic conductance from interneuron population *j* to pyramidal neuron *i* is denoted by 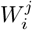, where *j* = *S, P* corresponds to the stimulus- and prediction-driven interneuron populations, respectively.

In the simulations, synapses originating from the same presynaptic interneuron population share a common gating variable, denoted by *s*_*j*_. The total inhibitory conductance received by a postsynaptic pyramidal neuron *i* from both interneuron populations is given by:

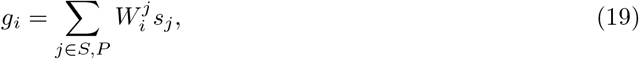

where *s*_*j*_ is the gating variable for population *j*.

Our model captures pyramidal neuron activity at both the single-cell and population levels. The network consists of *N*_neu_ pyramidal neurons, with two distinct interneuron populations relaying stimulus- and prediction-driven inhibition, respectively.

In the case where the homeostasis assumption is applied, the excitatory synaptic inputs from the stimulus and prediction are anti-correlated. The stimulus input strength is linearly spaced from a maximum value 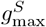 to zero. Thus, for the *i*-th neuron, the stimulus input strength is defined as:

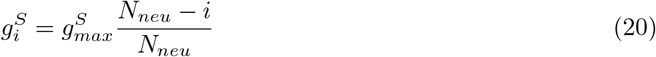

With input strength *u*_*S*_ *∈* [0, 1], the total excitatory current from the stimulus is:

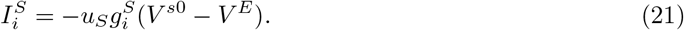

Similarly, the stimulus input to the *I*^*S*^ interneuron population is defined by the synaptic strength parameter 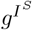. In the perturbation experiments, the perturbation level is normalized by this baseline input strength. A perturbation value of *I*^*S*^ perturb = −1 indicates that the *I*^*S*^ population receives no excitation when the stimulus is presented, whereas *I*^*S*^ perturb = 1 indicates that the *I*^*S*^ population receives double the baseline excitation.

The top-down input is modeled as a presynaptic predictor neuron firing at a rate of 5(1 + *u*_*P*_) Hz, where *u*_*P*_ *∈* [0, 1] represents the prediction strength. This yields a predictor firing rate in the range [5, 10] Hz. The synaptic weights from the predictor neuron to the dendritic compartments of pyramidal neurons are linearly spaced from zero to a maximum value 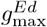. For the *i*-th neuron, the top-down connectivity strength is given by 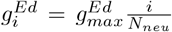. Furthermore, the excitatory synaptic input is composed of a mixture of NMDA and AMPA receptors, with a fraction *κ* mediated by NMDA synapses and the remaining (1 − *κ*) by AMPA synapses.

The input conductance is computed by substituting the predictor neuron’s firing rate into Equations 15 and 18. Specifically, the dynamics of the AMPA and NMDA gating variables become:

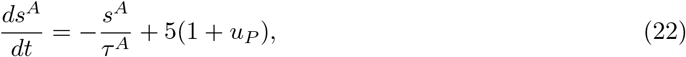

and

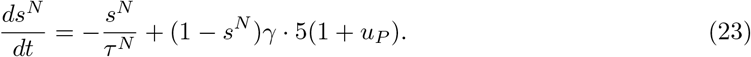

The input current to pyramidal neuron *i* depends on both the gating variables 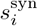 and the dendritic membrane potential 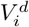:

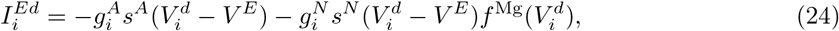

where 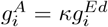 and 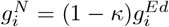.

Similarly, the prediction input to the *I*^*P*^ population is defined by the input strength 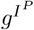. In the perturbation experiments, the degree of perturbation is normalized by the input strength to the *I*^*S*^ population, 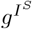, so that equivalent perturbation magnitudes are comparable across the *I*^*S*^ and *I*^*P*^ populations.

We further include background stimulus conductance to all neurons in the model, ensuring that each cell maintains a similar low, but non-zero, baseline firing rate.

In the realistic model without the homeostasis assumption, we shuffle the synaptic weights from the predictor neuron to pyramidal cells, such that the input strengths from the stimulus and prediction are uncorrelated.

We do not include lateral connectivity between pyramidal neurons in this model. While lateral inhibition may aid in differentiating responses between nPE and pPE neurons, it cannot contribute to canceling stimulus or prediction inputs in the expected condition, where activity is low across the population. For simplicity, we omit these connections from the model.

All parameters used in this study are listed in the Supplementary Table. All code will be available on GitHub upon publication acceptance.

### 4.6 Data analysis

In the motion-modulated auditory response analysis (Figure 3), we quantify the selective modulation effect of the prediction signal using the same modulation index (MI) as defined in [4]. If the prediction increases the response to the stimulus, the MI is positive; if it suppresses the response, the MI is negative. Specifically, for each cell, let *r*_*s*_ denote the response to the unmodulated stimulus and *r*_*m*_ the response to the modulated stimulus. We compute the angular deviation from the diagonal (indifference) line as: *θ* = arctan(*r*_*m*_*/r*_*s*_) − *π/*4 The resulting *θ ∈* [−*π, π*] is linearly rescaled to the range [−2, 2] for visualization clarity. An MI of −1 indicates that the response to the modulated stimulus is zero. Only cells with responses greater than 2 *×* (1 + 1.28), corresponding to a statistically significant response (*p* = 0.1) compared to the baseline rate *r* = 2, are included in the analysis.

The discriminability between stimulus pairs is computed to assess whether the sensory area can continue to process stimuli accurately. Discriminability is quantified using a form of the Fisher information metric [14], specifically the squared d-prime, summed across neurons:

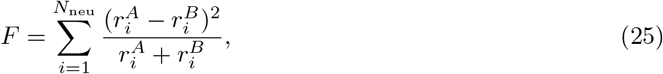

where *A* and *B* denote two different stimulus conditions. With this formulation, the Fisher information can be interpreted as the discriminability of an optimal linear decoder [40].

In the motion-vision mismatch test (Figure 4), we examine the emergence of a bimodal distribution in the correlation diagram following learning. To generate this distribution, we randomize the strengths of the stimulus and prediction inputs and run 200 trials continuously in the model. For each neuron, we calculate the correlation between its response and the input strengths *u*_*S*_ and *u*_*P*_. We then compute an angular difference, *θ*, in polar coordinates, where *θ* = 0 indicates equal correlation with stimulus and prediction. This angular metric is further weighted by the radial distance in the polar plane to emphasize neurons with larger response variance. A bimodal distribution of *θ* after learning reflects the emergence of both pPE and nPE neuronal populations. Because neuronal responses in the model are solely determined by the stimulus and prediction input strengths, the response variance is almost entirely explained by these two variables. Consequently, the data points lie approximately on a ring manifold in the correlation space.

We further apply the same analysis to data collected from C57BL/6J mice of either sex (*n* = 6 per group) under both coupled training (CT) and non-coupled training (UT) conditions, originally published in [2]. The total number of cells under examination is *N*_CT_ = 939 for the CT group, and *N*_NT_ = 690 for the NT group. We demonstrate that the bimodal distribution emerges only in the coupled training condition by fitting the angular data to Gaussian Mixture Models (GMMs) with one or two components [7].

To perform this analysis on weighted data, we first construct a pseudo-dataset. For each cell in polar coordinates with angle *θ*_*i*_ and radius *r*_*i*_, we replicate *θ*_*i*_ by 100 *×r*_*i*_ times to reflect the weighting. Only data with *θ*_*i*_ *∈* [−*π/*2, *π/*2] are included, in order to avoid spurious peaks near *θ* = −*π*. All replicated values are collected to form the pseudo-dataset. The resulting pseudo-dataset is then fitted to Gaussian Mixture Models using the MATLAB function *fitgmdist*, with either one or two components and default parameter settings.

To evaluate whether one-component or two-component fitting is superior, we use the Bayesian Information Criterion (BIC) [52]. Typically, a difference of ΔBIC *>* 10 is considered strong evidence in favor of the model with the lower BIC. In our analysis, fitting two-component GMMs to both pseudo CT and NT datasets yields ΔBIC *>* 10, favoring the two-component model in both cases (Figure 16). However, when we overlay the fitted GMMs on the pseudo-datasets, only the CT datasets show a true bimodal distribution. In contrast, the uncoupled condition shows a single dominant peak, suggesting that the means of the two Gaussian components are nearly identical (Figure 16A, blue dashed line).

## Code Availability

We analyze the model-generated data using Python 3.9 and the experimental data using MATLAB R2024b. All analysis code will be made available on GitHub upon acceptance.

## Declaration of generative AI and AI-assisted technologies in the writing process

During the preparation of this work, the authors used ChatGPT-4o in order to assist with proof-reading and improving the clarity of the manuscript. After using this tool, the authors reviewed and edited the content as needed and take full responsibility for the content of the publication.

## Author contributions

J.H.M. and X.J.W. conceptualized the project. J.H.M. performed simulations, data analyses, and wrote the manuscript. X.J.W. supervised the study.

## Acknowledgement

We thank Aldo Battista, Yue Liu, Tianshu Li, and other members of Xiao-Jing Wang lab for the discussion. We thank Cristina Savin and David Schneider for their suggestions and feedback on an early version of the manuscript. We thank Georg Keller for technical assistance regarding access to the publicly available dataset. This work is supported by ONR grant N00014-23-1-2040 and NIH grant R01MH062349 (to XJW)

## Supplementary Materials

### Local three-factor learning rule optimizes the mismatch calculation

In this section, we mathematically demonstrate that our local three-factor learning rule can optimize mismatch computation under a set of reasonable assumptions. We proceed in three parts: First, we show how different types of training samples influence a simplified optimization problem in a network with *N*_neu_ = 2, where only one *I*^*P*^ and one *I*^*S*^ population are present. Second, we prove that our three-factor learning rule converges to the same optimal solution as the gradient descent algorithm. Finally, we extend the analysis to the general case with *N*_neu_ *>* 2, in which neurons receive heterogeneous stimulus and prediction inputs. We conclude by specifying the conditions under which a neuron becomes a pPE or nPE unit.

### Gradient descent in different scenarios optimizes different weights

Here, we prove that when *N*_*neu*_ = 2, our learning algorithm is equivalent to solving the following optimization problem through gradient descent.

We assume a (or a pool of) neuron, denoted by *R*, represents a decoder that is optimized for the mismatch signal between prediction and stimulus. Also, we annotate the input *u*_*S*_, *u*_*P*_ directly as *S, P* for simplicity here. The cost function is defined as: :

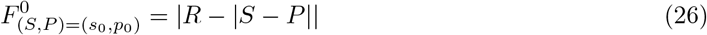

where *s*_0_, *p*_0_ represent a specific case. And we will minimize the integration of all the potential pairs (*s*_0_, *p*_0_)

The decoder neuron receives input from neurons in the sensory area. In the case of *N*_*neu*_ = 2, we further assume that one neuron receives strong stimulus input and another receives strong prediction input. There are two pools of interneurons that relay stimulus inhibition and prediction inhibition, respectively. These strong conditions will be released by heterogeneity later. The corresponding strength is 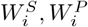 with *i* = 1, 2. For simplicity, we further assume that these neurons are activated through a ReLU function. Then, we will have

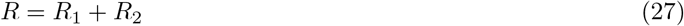

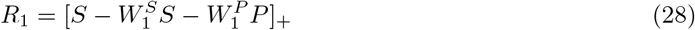

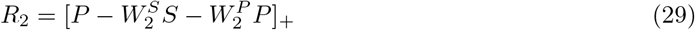

Then, the optimization problem can be written as

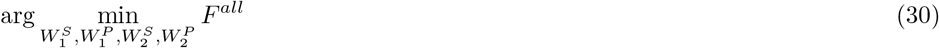

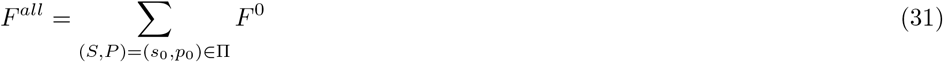

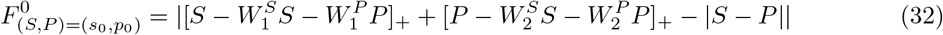

Obviously, the only optimal solution is 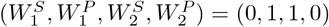.

Next, we will prove that the gradient descent method can achieve this only on the condition that the training set includes both expected samples and mismatched samples.

We first let *X* = *S* − *P* and *Y* = *S* + *P*, then we have

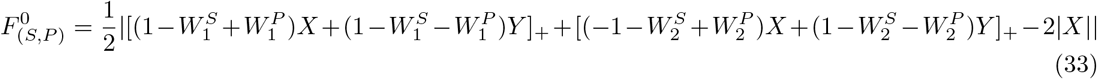

The training set is split into an expected set and a mismatch set. *F*^*all*^ = *F*^*E*^ + *F*^*MM*^ = ∑_|*P* − *S ≤ ϵ*|_ *F* ^0^ + ∑ _|*P* − *S* | *≥ϵ*_ *F* ^0^, while *ϵ* is a small amount.

We first consider the effect on training in the expected case, where the difference between stimulus and prediction is small (|*X*| = |*P* − *S*| *< ϵ*; *F*^*E*^), then:

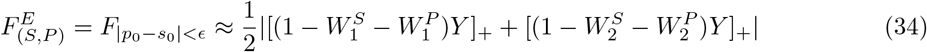

Since all the terms are non-negative, the optimal solution requires:

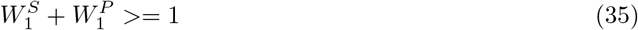

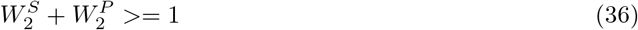

We first consider the initial condition satisfies 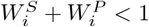 with *i* = 1, 2, an optimal solution

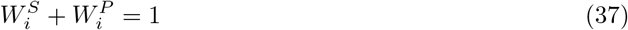

can be reached by updating the weight along the descending gradient:

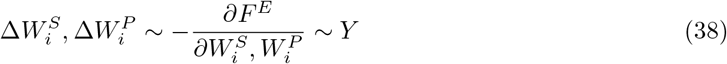

where *i* = 1, 2

The undetermined solution suggests that if we train this optimization problem only with the expected samples, the neuron will receive the same amount of inhibition as the excitation. Yet, the neuron cannot distinguish the source of inhibition. As a result, the optimal solution for mismatch calculation may not be reached.

We now consider the mismatch training set (|*X*| = |*P* − *S*| *> ϵ, F*^*MM*^). If we assume that the weight is already trained with sufficient expected cases, then the weights are constrained by 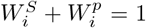 for *i* = 1, 2. Plug in equation 33, We have:

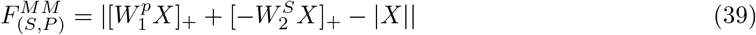

which gives the optimal solution

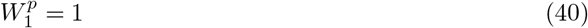

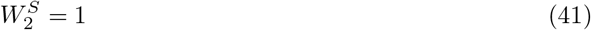

Reconsider the condition 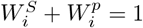, then

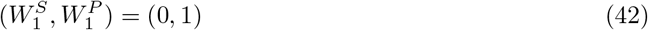

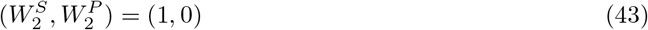

Notice 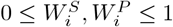, this optimal solution can be reached by gradient descent:

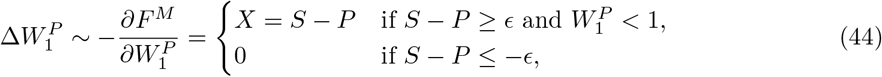

and 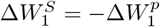.

Similarly,

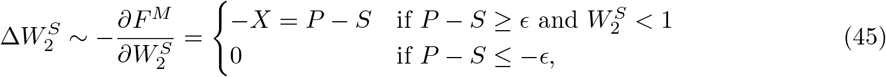

and 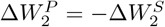

The update is illustrated in Figure 2B.

Now we consider the condition when 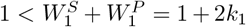. The case where the 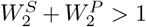 can be proven similarly. In this case, the synaptic weight 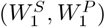 won’t be updated in the expected set *F*^*E*^ since the gradient is zero. In the mismatched case *F*^*MM*^ with *X >* 0, we will have

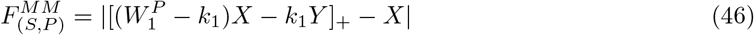

Since *X, Y >* 0, we will have

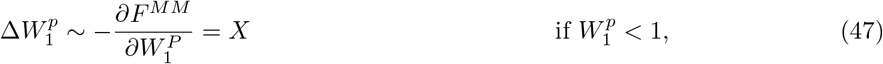

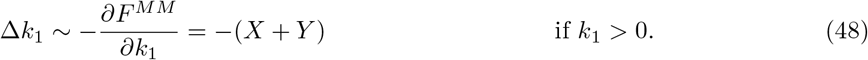

The learning will stops at 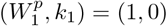. Since 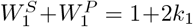, this suggests that eventually the solution will be the same 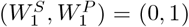. Again, here training only in the expected case *F*^*E*^ cannot generate a unique solution.

#### 4.6.1 The three-factor learning rule can optimize the mismatch calculation as the gradient descent

In this section, we will prove that our simple learning rule can achieve the optimized mismatch calculation.

Our three-factor learning rule satisfies

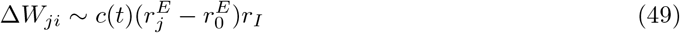

Our *c*(*t*) = 2(*c*_0_ −| *S* − *P*|) with *c*_0_ is a constant, 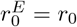 is a small target firing rate. The *c*(*t*) here represents neuromodulator activity, like noradrenalin [32, 30].

We first show the behavior of learning on 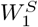 and 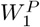 with the initial condition 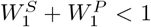 and learning within the expected set (|*S* − *P*| *< c*_0_; *F*^*E*^). In this case, *c*(*t*) *>* 0, and:

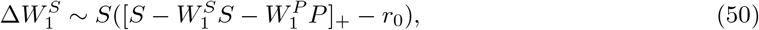

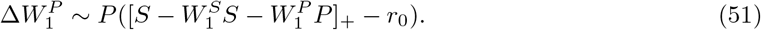

Since *S ≈ P*, this leads to an update 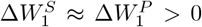. Also, *r*_0_ is small, the learning stops at 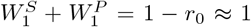, leading to a family of solutions instead of a unique one, similar to the case of gradient descent within *F*^*E*^ (Figure 2C). We will refer to the line 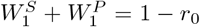 as the slow learning manifold in the following derivations. Also, if the initial condition starts with excessive inhibition 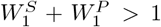, the nonzero targeted firing rate *r*_0_ *>* 0 leads to 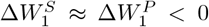, converging to the same slow learning manifold 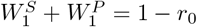.

When learning within the mismatched set *F*^*MM*^ : |*S* − *P*| *> c*_0_, we have *c*(*t*) *<* 0. Then

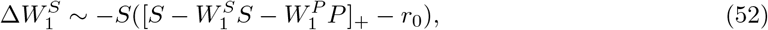

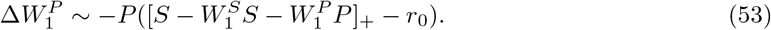

Only when *S* − *P > c*_0_, the term in the rectifier is positive, such that the weights 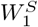 and 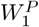 are updated. In contrast, when *S* − *P <* −*c*_0_, the weights *W*^*S*^ and *W*^*P*^ are not updated. Furthermore, when *S* − *P > c*_0_, we have *S > P* + *c*_0_ *> P*, then

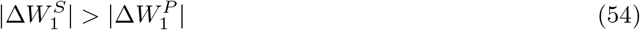

If the mismatch case is scarce, then the connectivity weight is constrained on the slow learning manifold. As a result, we have 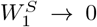 and 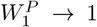 through a zigzag in the phase space (Figure 2C).

Since the exact value of *c*(*t*) only changes the learning speed, the conclusion still holds if we use an estimated piece-wise third factor 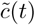, as shown in Supp. Figure 12.

#### 4.6.2 Behavior of the three-factor learning rule in a generalized case

In reality, both pyramidal cells and interneurons receive both top-down prediction signals and bottom-up stimulus signals. Without losing generality, we assume the following.

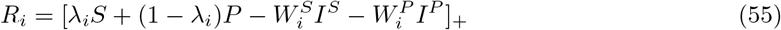

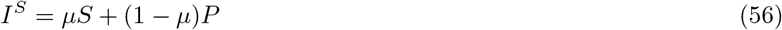

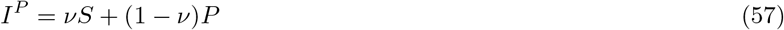

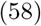

with neuron index *i*. Further assume *µ >* 0.5, and *ν <* 0.5, such that *I*_1_ receives stronger input from stimulus input than from prediction, and vice versa for *I*_2_. We drop the subindex *i* in the following section.

Again, within the expected cases *F*^*E*^, we have

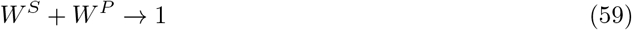

So we assume the initial weight **W**(*t*_1_) = (*W*^*S*^(*t*_1_), *W*^*P*^ (*t*_1_)) satisfies *W*^*S*^(*t*_1_) + *W*^*P*^ (*t*_1_) = 1.

And within the mismatched cases *F*^*MM*^, we have

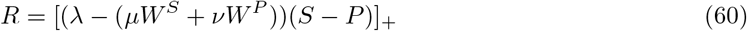

When *S > P* + *c*_0_ *> P*, if the initial condition satisfies *λ*− (*µW*^*S*^ + *νW*^*P*^) *>* 0, then *R >* 0. Further *µ >* 0.5 *> ν*, so we have *I*^*S*^ *> I*^*P*^. In addition, *c*(*t*) *<* 0 in the *F*^*MM*^. Plug everything into Equation49, we have

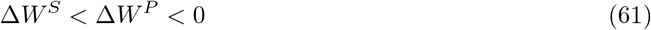

So the update once in the mismatches case is still the same as in Figure 2C.

To make sure the learning continues, we further check whether the learning process breaks the condition *λ* − (*µW*^*S*^ + *νW*^*P*^) *>* 0. In this scenario, we consider the update in the mismatched training case *F*^*MM*^, and the weight falls back to **W**(*t*_2_) = (*W*^*S*^(*t*_2_), *W*^*P*^ (*t*_2_)) on the slow learning manifold by the expected training cases *F*^*E*^. Then assume Δ*w*_0_ = *W*^*P*^ (*t*_2_) − *W*^*P*^ (*t*_1_) = −(*W*^*S*^(*t*_2_) − *W*^*S*^(*t*_1_)) *>* 0

Then

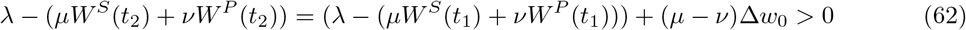

Since every term here is positive, the condition *λ* − (*µW*^*S*^ + *νW*^*P*^) *>* 0 is satisfied.

As a result, the learning process continues and eventually converges on the optimal solution (*W*^*S*^, *W*^*P*^) = (0, 1)

The case with *λ* − (*µW*^*S*^ + *νW*^*P*^) *<* 0 is similar.

To summarize, our learning rule will lead to

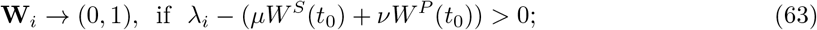

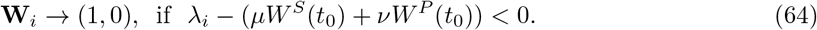

or in a more intuitive way

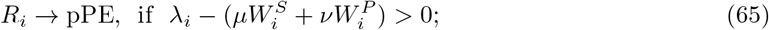

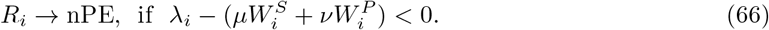

Whether a neuron becomes labeled as a pPE or nPE unit after learning is determined by two factors. First, if the neuron’s initial excitatory input is stronger from the stimulus than from the prediction, it is more likely to develop into a pPE neuron, and *vice versa*. Second, if the initial local inhibition is biased toward stimulus-driven inhibition, more neurons are likely to become nPE neurons, and *vice versa*.

One caveat is that this learning rule can lead to binary connectivity from the *I*^*S*^ and *I*^*P*^ populations to the pyramidal neurons. It is important to note that this outcome also arises when synaptic weights are updated via gradient descent. Although such binary configurations are theoretically possible, we find little empirical support for them in the literature. Therefore, we introduce an additional regularization mechanism to halt learning as follows:

One caveat point is that this will lead to a binary connection from *I*^*S*^, *I*^*P*^ to the pyramidal population. It is worth noticing that this is also true if we update through gradient descent. Even though this is possible, we find scarce literature supporting it. So we impose one additional regulator as follows to stop learning:

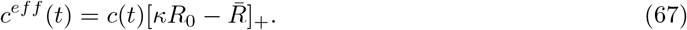

This condition applies when *c*(*t*) *<* 0. Here, 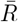 is the averaged firing rate of the pyramidal neurons, *R*_0_ is the small targeted firing rate as defined previously, *κ* is a scaling factor that determines the threshold beyond which further learning is suppressed.

This condition represents a naive downstream decoder that receives unbiased input from the *R*_*i*_ neurons and interrupts the local learning when its activity becomes sufficiently large. As a result, the final distance between the learned weights 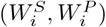 and the optimal solution, either (1, 0) or (0, 1), is determined by the initial bias in neuronal activity toward either the stimulus or prediction input.

For example, for two neurons *i, j* that both satisfy 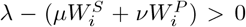, when *P > S*, the only term in Equation 49 that differentiates these two neurons is the firing rate *R*. I.e., the relative updating amplitude is determined by the firing rate of these two neurons

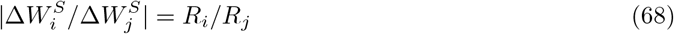

A proper choice *κ* will lead to a graded distribution of synaptic weight *W*^*S*^ and *W*^*P*^, as we observed in the simulation for both ReLU neurons (Supp. Figure 9) or realistic neurons (Figure 3, Supp. Figure 11).

## References

[1] Larry F Abbott and Frances S Chance. “Drivers and modulators from push-pull and balanced synaptic input”. In: Progress in brain research 149 (2005), pp. 147–155.

[2] Alexander Attinger, Bo Wang, and Georg B Keller. “Visuomotor coupling shapes the functional development of mouse visual cortex”. In: Cell 169.7 (2017), pp. 1291–1302.

[3] Nicholas J Audette and David M Schneider. “Stimulus-specific prediction error neurons in mouse auditory cortex”. In: Journal of Neuroscience 43.43 (2023), pp. 7119–7129.

[4] Nicholas J Audette et al. “Precise movement-based predictions in the mouse auditory cortex”. In: Current Biology 32.22 (2022), pp. 4925–4940.

[5] Danielle S Bassett et al. “Learning-induced autonomy of sensorimotor systems”. In: Nature neuroscience 18.5 (2015), pp. 744–751.

[6] Guillaume Bellec et al. “A solution to the learning dilemma for recurrent networks of spiking neurons”. In: Nature communications 11.1 (2020), p. 3625.

[7] Christopher M Bishop and Nasser M Nasrabadi. Pattern recognition and machine learning. Vol. 4. 4. Springer, 2006.

[8] Stephanie Bissière, Yann Humeau, and Andreas Lüthi. “Dopamine gates LTP induction in lateral amygdala by suppressing feedforward inhibition”. In: Nature neuroscience 6.6 (2003), pp. 587–592.

[9] Nicole K Bolt and Janeen D Loehr. “The auditory P2 differentiates self-from partner-produced sounds during joint action: contributions of self-specific attenuation and temporal orienting of attention”. In: Neuropsychologia 182 (2023), p. 108526.

[10] Zuzanna Brzosko, Susanna B Mierau, and Ole Paulsen. “Neuromodulation of spike-timing-dependent plasticity: past, present, and future”. In: Neuron 103.4 (2019), pp. 563–581.

[11] Chiayu Q Chiu, Andrea Barberis, and Michael J Higley. “Preserving the balance: diverse forms of long-term GABAergic synaptic plasticity”. In: Nature Reviews Neuroscience 20.5 (2019), pp. 272–281.

[12] Cameron Condylis et al. “Dense functional and molecular readout of a circuit hub in sensory cortex”. In: Science 375.6576 (2022), eabl5981.

[13] Phil R Corlett et al. “Toward a neurobiology of delusions”. In: Progress in neurobiology 92.3 (2010), pp. 345–369.

[14] Thomas M Cover. Elements of information theory. John Wiley & Sons, 1999.

[15] Trinity B Crapse and Marc A Sommer. “Corollary discharge across the animal kingdom”. In: Nature Reviews Neuroscience 9.8 (2008), pp. 587–600.

[16] CPJ De Kock et al. “Layer-and cell-type-specific suprathreshold stimulus representation in rat primary somatosensory cortex”. In: The Journal of physiology 581.1 (2007), pp. 139–154.

[17] Shlomo Dellal et al. “Inhibitory and disinhibitory VIP IN-mediated circuits in neocortex”. In: bioRxiv (2025).

[18] Steven J Eliades and Xiaoqin Wang. “Neural substrates of vocalization feedback monitoring in primate auditory cortex”. In: Nature 453.7198 (2008), pp. 1102–1106.

[19] Tatiana A Engel and Xiao-Jing Wang. “Same or different? A neural circuit mechanism of similarity-based pattern match decision making”. In: Journal of Neuroscience 31.19 (2011), pp. 6982–6996.

[20] Brandon J Farley et al. “Stimulus-specific adaptation in auditory cortex is an NMDA-independent process distinct from the sensory novelty encoded by the mismatch negativity”. In: Journal of Neuroscience 30.49 (2010), pp. 16475–16484.

[21] Nicolas Frémaux and Wulfram Gerstner. “Neuromodulated spike-timing-dependent plasticity, and theory of three-factor learning rules”. In: Frontiers in neural circuits 9 (2016), p. 85.

[22] Chris Frith. “The neural basis of hallucinations and delusions”. In: Comptes Rendus. Biologies 328.2 (2005), pp. 169–175.

[23] Aleena R Garner and Georg B Keller. “A cortical circuit for audio-visual predictions”. In: Nature neuroscience 25.1 (2022), pp. 98–105.

[24] Wulfram Gerstner et al. “Eligibility traces and plasticity on behavioral time scales: experimental support of neohebbian three-factor learning rules”. In: Frontiers in neural circuits 12 (2018), p. 53.

[25] Nathan W Gouwens et al. “Integrated morphoelectric and transcriptomic classification of cortical GABAergic cells”. In: Cell 183.4 (2020), pp. 935–953.

[26] Manu Srinath Halvagal and Friedemann Zenke. “The combination of Hebbian and predictive plasticity learns invariant object representations in deep sensory networks”. In: Nature Neuroscience 26.11 (2023), pp. 1906–1915.

[27] Donald Olding Hebb. The organization of behavior: A neuropsychological theory. Psychology press, 2005.

[28] Loreen Hertäg and Claudia Clopath. “Prediction-error neurons in circuits with multiple neuron types: Formation, refinement, and functional implications”. In: Proceedings of the National Academy of Sciences 119.13 (2022), e2115699119.

[29] Loreen Hertäg and Henning Sprekeler. “Learning prediction error neurons in a canonical interneuron circuit”. In: Elife 9 (2020), e57541.

[30] Rebecca Jordan. “The locus coeruleus as a global model failure system”. In: Trends in Neurosciences 47.2 (2024), pp. 92–105.

[31] Rebecca Jordan and Georg B Keller. “Opposing influence of top-down and bottom-up input on excitatory layer 2/3 neurons in mouse primary visual cortex”. In: Neuron 108.6 (2020), pp. 1194–1206.

[32] Rebecca Jordan and Georg B Keller. “The locus coeruleus broadcasts prediction errors across the cortex to promote sensorimotor plasticity”. In: Elife 12 (2023), RP85111.

[33] Mahesh M Karnani, Masakazu Agetsuma, and Rafael Yuste. “A blanket of inhibition: functional inferences from dense inhibitory connectivity”. In: Current opinion in Neurobiology 26 (2014), pp. 96–102.

[34] Georg B Keller and Thomas D Mrsic-Flogel. “Predictive processing: a canonical cortical computation”. In: Neuron 100.2 (2018), pp. 424–435.

[35] John H Krystal et al. “NMDA receptor antagonist effects, cortical glutamatergic function, and schizophrenia: toward a paradigm shift in medication development”. In: Psychopharmacology 169 (2003), pp. 215–233.

[36] Dimitri M Kullmann et al. “Plasticity of inhibition”. In: Neuron 75.6 (2012), pp. 951–962.

[37] Lukasz Kuśmierz, Takuya Isomura, and Taro Toyoizumi. “Learning with three factors: modulating Hebbian plasticity with errors”. In: Current opinion in neurobiology 46 (2017), pp. 170–177.

[38] Markus Lappe, Frank Bremmer, and Albert V van den Berg. “Perception of self-motion from visual flow”. In: Trends in cognitive sciences 3.9 (1999), pp. 329–336.

[39] Matthew E Larkum, Walter Senn, and Hans-R Lüscher. “Top-down dendritic input increases the gain of layer 5 pyramidal neurons”. In: Cerebral cortex 14.10 (2004), pp. 1059–1070.

[40] John Hongyu Meng and Hermann Riecke. “Structural spine plasticity: Learning and forgetting of odor-specific subnetworks in the olfactory bulb”. In: PLoS computational biology 18.10 (2022), e1010338.

[41] Martial Mermillod, Aurélia Bugaiska, and Patrick Bonin. The stability-plasticity dilemma: Investigating the continuum from catastrophic forgetting to age-limited learning effects. 2013.

[42] Cristopher M Niell and Michael P Stryker. “Highly selective receptive fields in mouse visual cortex”. In: Journal of Neuroscience 28.30 (2008), pp. 7520–7536.

[43] Sean M O’Toole, Hassana K Oyibo, and Georg B Keller. “Molecularly targetable cell types in mouse visual cortex have distinguishable prediction error responses”. In: Neuron 111.18 (2023), pp. 2918–2928.

[44] Anne-Marie M Oswald and Alex D Reyes. “Maturation of intrinsic and synaptic properties of layer 2/3 pyramidal neurons in mouse auditory cortex”. In: Journal of neurophysiology 99.6 (2008), pp. 2998–3008.

[45] Verena Pawlak et al. “Timing is not everything: neuromodulation opens the STDP gate”. In: Frontiers in synaptic neuroscience 2 (2010), p. 146.

[46] Carsten K Pfeffer et al. “Inhibition of inhibition in visual cortex: the logic of connections between molecularly distinct interneurons”. In: Nature neuroscience 16.8 (2013), pp. 1068–1076.

[47] Rajesh PN Rao and Dana H Ballard. “Predictive coding in the visual cortex: a functional interpretation of some extra-classical receptive-field effects”. In: Nature neuroscience 2.1 (1999), pp. 79–87.

[48] Alfonso Renart, Pengcheng Song, and Xiao-Jing Wang. “Robust spatial working memory through homeostatic synaptic scaling in heterogeneous cortical networks”. In: Neuron 38.3 (2003), pp. 473–485.

[49] Shuzo Sakata and Kenneth D Harris. “Laminar structure of spontaneous and sensory-evoked population activity in auditory cortex”. In: Neuron 64.3 (2009), pp. 404–418.

[50] David M Schneider, Anders Nelson, and Richard Mooney. “A synaptic and circuit basis for corollary discharge in the auditory cortex”. In: Nature 513.7517 (2014), pp. 189–194.

[51] David M Schneider, Janani Sundararajan, and Richard Mooney. “A cortical filter that learns to suppress the acoustic consequences of movement”. In: Nature 561.7723 (2018), pp. 391–395.

[52] Gideon Schwarz. “Estimating the dimension of a model”. In: The annals of statistics (1978), pp. 461–464.

[53] Geun Hee Seol et al. “Neuromodulators control the polarity of spike-timing-dependent synaptic plasticity”. In: Neuron 55.6 (2007), pp. 919–929.

[54] Magdalena Solyga and Georg B Keller. “Multimodal mismatch responses in mouse auditory cortex”. In: eLife 13 (2025), RP95398.

[55] Marc A Sommer and Robert H Wurtz. “A pathway in primate brain for internal monitoring of movements”. In: science 296.5572 (2002), pp. 1480–1482.

[56] Marc A Sommer and Robert H Wurtz. “Brain circuits for the internal monitoring of movements”. In: Annu. Rev. Neurosci. 31.1 (2008), pp. 317–338.

[57] John Sweller. “Cognitive load during problem solving: Effects on learning”. In: Cognitive science 12.2 (1988), pp. 257–285.

[58] John Sweller. “Cognitive load theory”. In: Psychology of learning and motivation. Vol. 55. Elsevier, 2011, pp. 37–76.

[59] Nevo Taaseh, Amit Yaron, and Israel Nelken. “Stimulus-specific adaptation and deviance detection in the rat auditory cortex”. In: PloS one 6.8 (2011), e23369.

[60] Tim P Vogels et al. “Inhibitory plasticity balances excitation and inhibition in sensory pathways and memory networks”. In: Science 334.6062 (2011), pp. 1569–1573.

[61] Felix C Widmer, Sean M O’Toole, and Georg B Keller. “NMDA receptors in visual cortex are necessary for normal visuomotor integration and skill learning”. In: Elife 11 (2022), e71476.

[62] Kong-Fatt Wong and Xiao-Jing Wang. “A recurrent network mechanism of time integration in perceptual decisions”. In: Journal of Neuroscience 26.4 (2006), pp. 1314–1328.

[63] Baba Yogesh and Georg B Keller. “Activity in serotonergic axons in visuomotor areas of cortex is modulated by the recent history of visuomotor coupling”. In: bioRxiv (2025), pp. 2025–03.

[64] Friedemann Zenke and Wulfram Gerstner. “Hebbian plasticity requires compensatory processes on multiple timescales”. In: Philosophical transactions of the royal society B: biological sciences 372.1715 (2017), p. 20160259.

[65] Petr Znamenskiy et al. “Functional specificity of recurrent inhibition in visual cortex”. In: Neuron 112.6 (2024), pp. 991–1000.

